# An atlas of healthy and injured cell states and niches in the human kidney

**DOI:** 10.1101/2021.07.28.454201

**Authors:** Blue B. Lake, Rajasree Menon, Seth Winfree, Qiwen Hu, Ricardo Melo Ferreira, Kian Kalhor, Daria Barwinska, Edgar A. Otto, Michael Ferkowicz, Dinh Diep, Nongluk Plongthongkum, Amanda Knoten, Sarah Urata, Abhijit S. Naik, Sean Eddy, Bo Zhang, Yan Wu, Diane Salamon, James C. Williams, Xin Wang, Karol S. Balderrama, Paul Hoover, Evan Murray, Anitha Vijayan, Fei Chen, Sushrut S. Waikar, Sylvia Rosas, Francis P. Wilson, Paul M. Palevsky, Krzysztof Kiryluk, John R. Sedor, Robert D. Toto, Chirag Parikh, Eric H. Kim, Evan Z. Macosko, Peter V. Kharchenko, Joseph P. Gaut, Jeffrey B. Hodgin, Michael T. Eadon, Pierre C. Dagher, Tarek M. El-Achkar, Kun Zhang, Matthias Kretzler, Sanjay Jain, for the KPMP consortium

## Abstract

Understanding kidney disease relies upon defining the complexity of cell types and states, their associated molecular profiles, and interactions within tissue neighborhoods. We have applied multiple single-cell or -nucleus assays (>400,000 nuclei/cells) and spatial imaging technologies to a broad spectrum of healthy reference (n = 42) and disease (n = 42) kidneys. This has provided a high resolution cellular atlas of 100 cell types that include rare and novel cell populations. The multi-omic approach provides detailed transcriptomic profiles, epigenomic regulatory factors, and spatial localizations for major cell types spanning the entire kidney. We further identify and define cellular states altered in kidney injury, encompassing cycling, adaptive or maladaptive repair, transitioning and degenerative states affecting several segments. Molecular signatures of these states permitted their localization within injury neighborhoods using spatial transcriptomics, and large-scale 3D imaging analysis of ∼1.2 million neighborhoods provided linkages to active immune responses. These analyses further defined biological pathways relevant to injury niches, including signatures underlying the transition from reference to predicted maladaptive states that were associated with a decline in kidney function during chronic kidney disease. This human kidney cell atlas, including injury cell states and neighborhoods, will be a valuable resource for future studies.

## Introduction

The human kidneys play vital systemic roles in the preservation of body fluid homeostasis, metabolic waste product removal and blood pressure maintenance. This organ system has a remarkable ability to perform its functions by adapting to a wide range of physiological demands and pathological insults. After injury, there are dynamic acute and chronic morphological and cellular changes in renal tubules and surrounding interstitial niche. The balance between successful or maladaptive repair processes may ultimately determine potential for progressive decline in kidney function over time^1–4^. In this regard it is critical to delineate the landscape of cellular and molecular diversity of gene expression and regulation at a single cell level in the human kidney. This will be needed to fully understand how acute kidney injury (AKI) events can increase risk for progression to chronic kidney disease (CKD), kidney failure, heart disease or death, issues that remain a global concern^5, 6^.

To this end, we report a next-generation multimodal single cell and 3D atlas that leverages integrated transcriptomic, epigenomic and imaging data over three major consortia: the Human Biomolecular Atlas Program (HuBMAP)^7^, the Kidney Precision Medicine Project (KPMP)^8^, and the Human Cell Atlas (HCA)^9^. To ensure robust cell state profiles, reference tissues were obtained from multiple sources, and biopsies were collected from AKI and CKD patients under rigorous quality assurance and control procedures^7, 8, 10^. We define micro niches for healthy and altered states across different regions of the human kidney spanning the cortex and medulla, to the papillary tip, and identify gene expression and regulatory modules in altered states associated with worsening kidney function. The resultant atlas of molecular cell types and their spatially resolved healthy and injury niches greatly expands upon existing efforts^11–14^. This will serve as an important resource for a broad user base of investigators and clinicians working towards a better understanding of kidney processes in health or disease.

## Results

### Constructing a Cellular Atlas of the Human Kidney

To fully interrogate molecularly defined kidney cell types, we have applied droplet-based transcriptomic assays (Chromium v3) for single nuclei (snCv3) and single cells (scCv3) and the dual transcriptomic/epigenomic assay for single-nucleus chromatin accessibility and mRNA expression sequencing (SNARE-seq2 or SNARE2)^15, 16^ to a broad range of tissues from reference to AKI and CKD biopsies (**Supplementary Tables 1-3**). To glean insights into biologically relevant spatial interactions between these cell types or states *in situ*, we further applied 3D label-free imaging, multiplex fluorescence imaging, and the spatial transcriptomic assays Slide-Seq2^17, 18^ and Visium (**Fig. 1, Supplementary Tables 1-2; Methods**). Our heterogeneous sampling approach was designed to ensure cell type discovery with minimal assay dependent biases or artifacts associated with different sources of reference or disease kidney samples.

**Figure 1.**
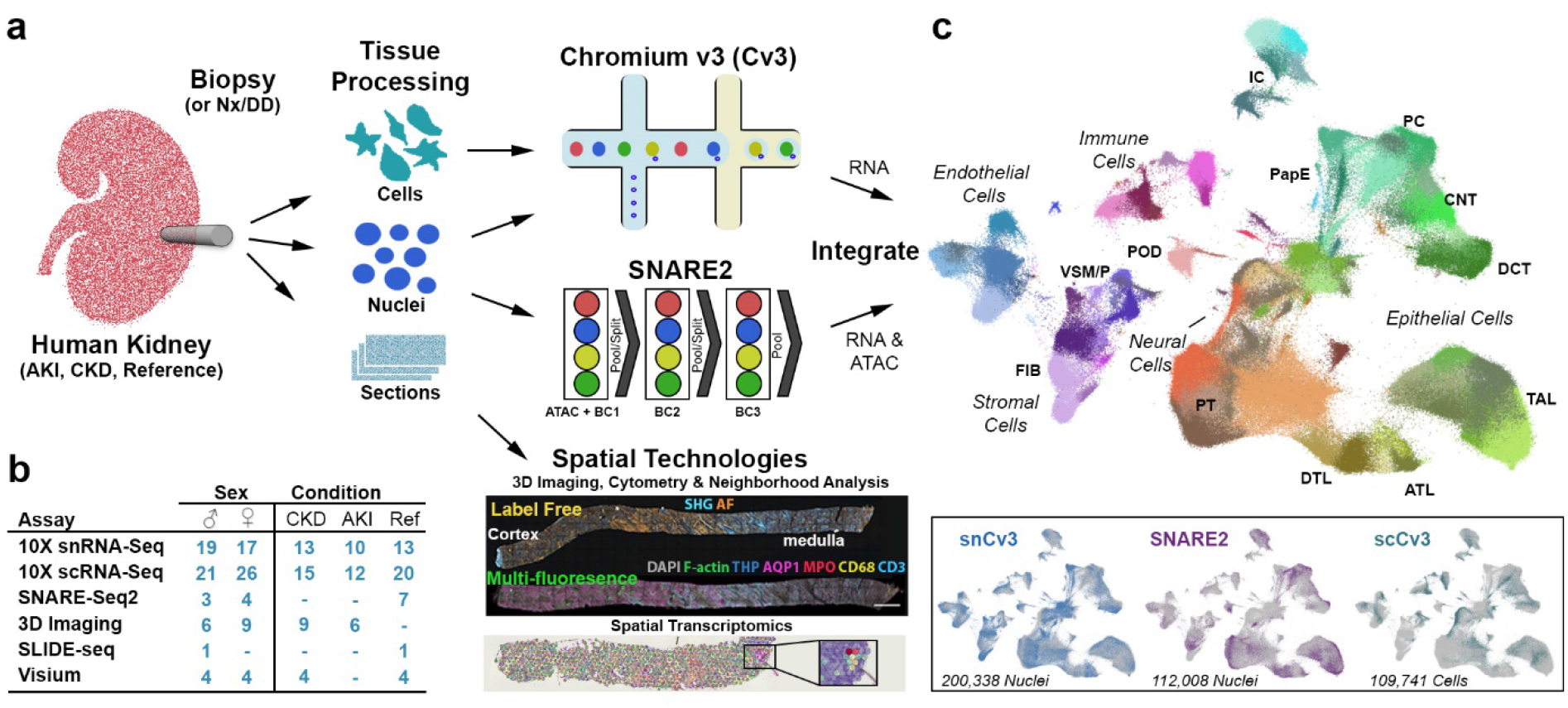
Overview of technologies used to generate a human kidney cell atlas. **a.** Human kidney samples summarized in (**b**) consisted of healthy reference, AKI or CKD nephrectomies (Nx), deceased donors (DD) or biopsies. Tissues were processed for one or more assays that included snCv3, scCv3, SNARE2, 3D imaging or spatial transcriptomics (Slide-seq2, Visium). **c.** Omic RNA data was integrated, as shown by joint UMAP embedding, for alignment of cell type annotations across the three different data modalities.

Integrative cross-platform transcriptome analyses were performed on >400,000 nuclei/cells (after quality filtering, **Methods**) from 58 reference tissues (37 donors) and 52 diseased tissues (36 patients) that covered the spectrum of kidney health through to acute and chronic kidney disease (**Fig. 1, Extended Data Fig. 1**-**4, Supplementary Table 1**). Unsupervised clustering was first performed on snCv3, permitting discovery of 100 distinct cell populations, which were annotated to subclasses of epithelial, endothelial, stromal, immune and neural cell types (**Fig. 2**, **Extended Data Fig. 1-2, Supplementary Tables 4-5, Methods**). To further extend cell type annotations across omic platforms, snCv3 was used to anchor scCv3 (**Extended Data Fig. 3**) and SNARE2 (**Extended Data Fig. 4**) data sets to the same embedding space, and cell type labels were assigned through integrative clustering (**Supplementary Tables 6-7, Methods**). This permitted a single harmonized annotation across technologies for more accurate cross-platform interrogation of the same cell populations (**Extended Data Fig. 3**-**4**). This combined omic atlas permitted deeper and cross-validated molecular profiles for these aligned kidney cell types, leveraging the distinct advantages of each technology, for instance the addition of cytosolic transcripts from scCv3 and regulatory elements from SNARE2 accessible chromatin (AC).

**Figure 2.**
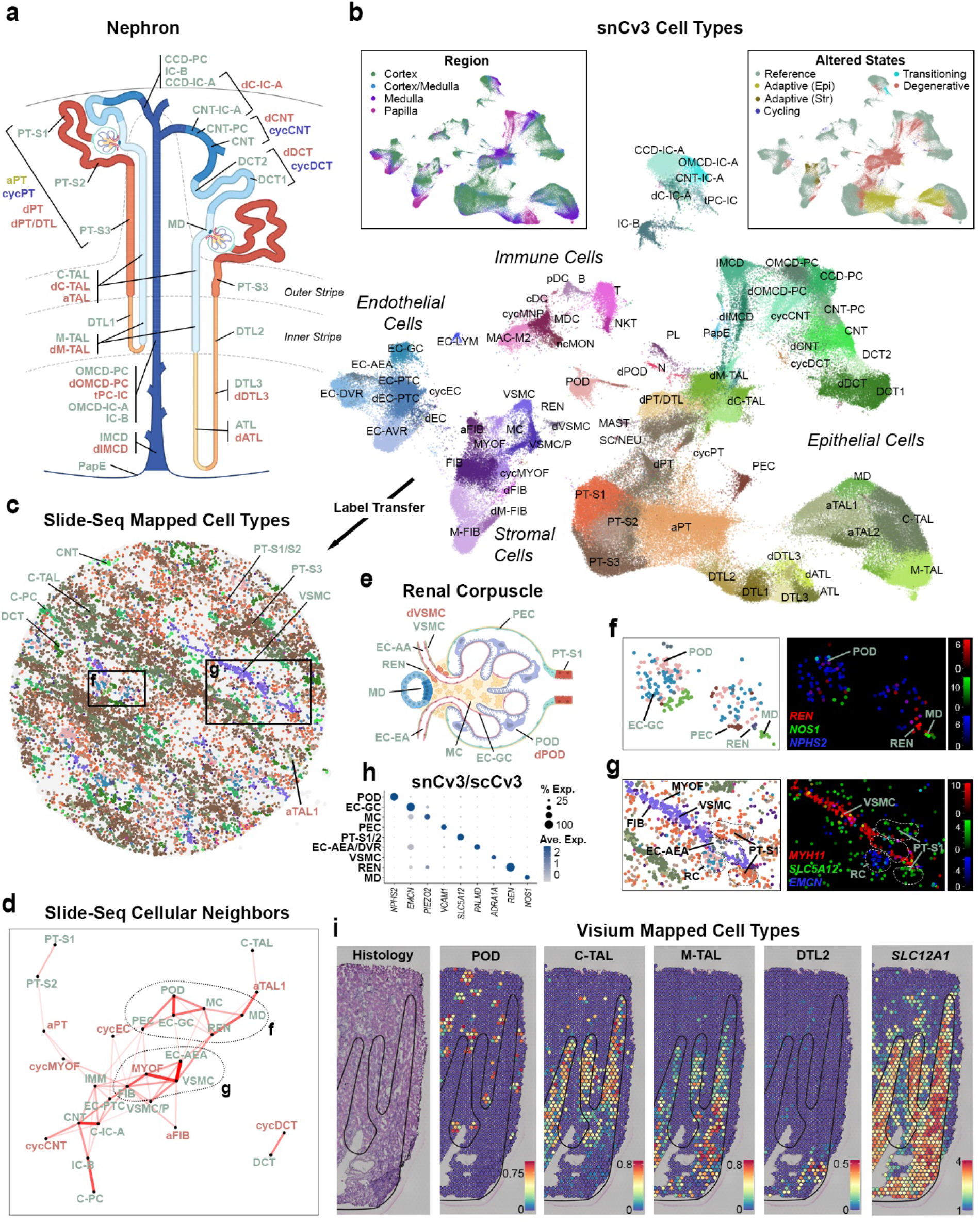
Spatially resolved atlas of molecular cell types. **a.** Schematic of the human nephron showing cell types and states resolved from snCv3. **b.** UMAP embedding showing cell types (subclass level 3) for snCv3. Insets show overlays for both regional origin and altered state status. **c.** Spatial distribution of cell type labeled beads associated with a single Slide-seq2 processed tissue puck. Puck diameter is 3mm. **d.** Cell proximity network for Slide-seq2 cell types. **e.** Schematic of the renal corpuscle showing snCv3 resolved cell types. **f.** Left panel shows Slide-seq2 puck area indicated in (**c**) and predicted cell types for renal corpuscles, highlighting cellular neighbors predicted in (**d**). Right panel shows the mapped expression values for corresponding marker genes. **g.** Left panel shows Slide-seq2 puck area indicated in (**c**) and predicted cell types for the AEAs and surrounding cell types, highlighting cellular neighbors predicted in (**d**). Right panel shows the mapped expression values for corresponding marker genes. **h.** Dotplot showing average expression values in snCv3 and scCv3 for markers shown in (**f**) and (**g**). **i.** 10X Visium data on a healthy reference kidney (cortex, top; medulla, bottom). Left panel shows H&E staining of the tissue, right panels show per bead predicted transfer scores for cell clusters or transcript expression values. Each spot is 55 μm in diameter.

**Figure 3.**
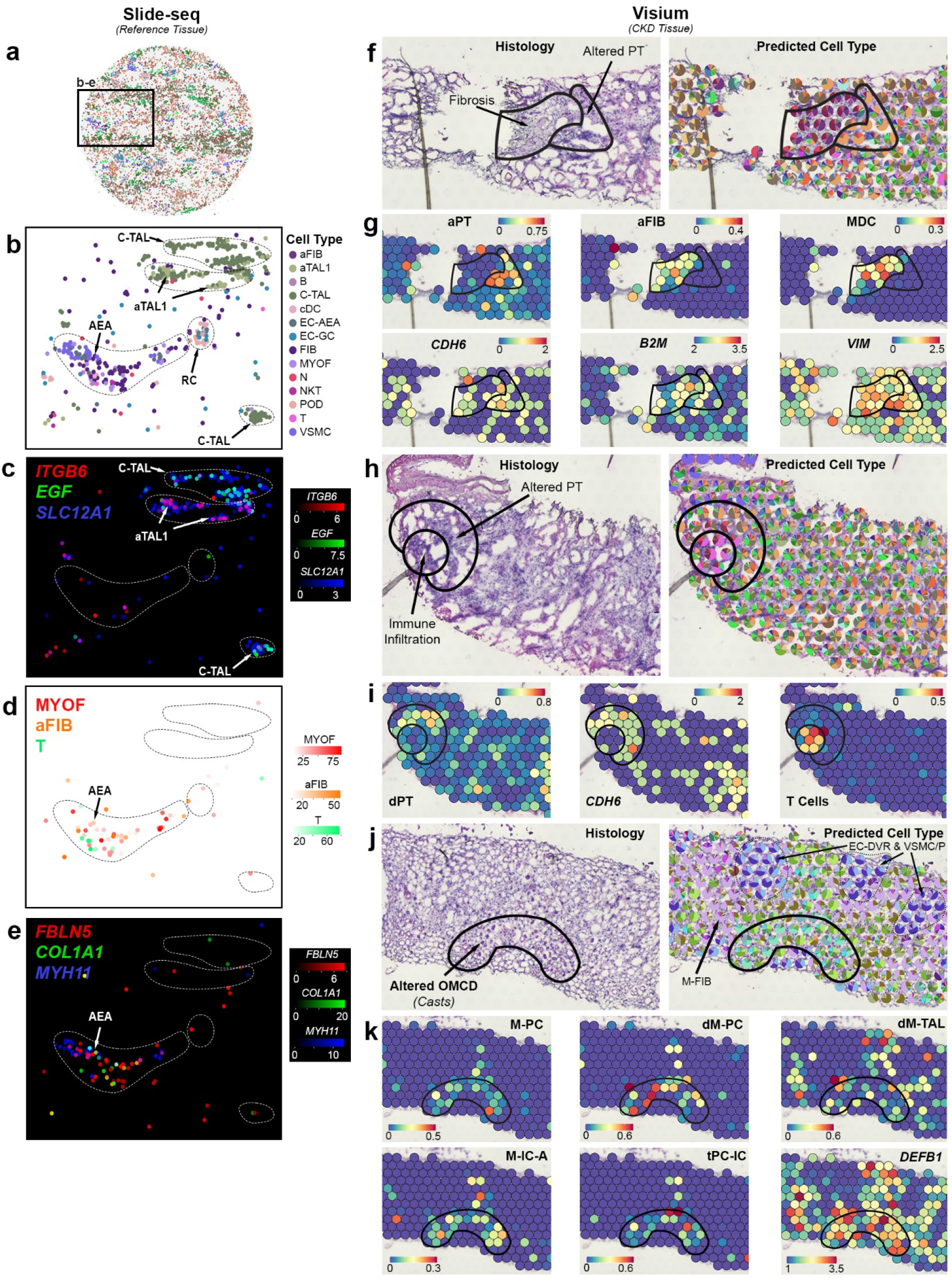
Transcriptomically defined injury neighborhoods. **a.** Slide-seq2 bead locations for a single tissue puck, colored by predicted cell subclasses as shown in Figure 2b. Puck diameter is 3mm. **b.** Slide-seq2 puck region indicated in (**a**) showing a subset of predicted cell types. **c.** Mapped expression values for aTAL (*ITGB6*) and TAL (*EGF* and *SLC12A1*) marker genes for cell types shown in (**b**). **d.** Prediction weights for cell types mapped to puck region indicated in (**a**). **e.** Mapped expression values for FIB (*FBLN5*), VSMC and MYOF (*MYH11*) and aStr (*COL1A1*) marker genes for cell types shown in (**d**). **f.** Histology and predicted cell types in a cortical region (CKD) of interstitial fibrosis. Pie charts are proportions of predicted transfer scores. **g.** Per bead predicted transfer scores for cell types or transcript expression values for area shown in (**f**). **h.** Histology and predicted cell types for a region with altered PT and immune cell infiltration. **i.** Predicted transfer scores and expression transcript expression values for area shown in (**h**). **j.** Histology and predicted cell types for a medullary region of acute tubular necrosis (cellular cast formation within tubular lumens, loss of brush border, loss of nuclei, and epithelial simplification). Pie charts are proportions of predicted transfer scores. **k.** Predicted transfer scores and expression transcript expression values for area shown in (**j**). For Visium panels, each spot is 55 μm in diameter.

**Figure 4.**
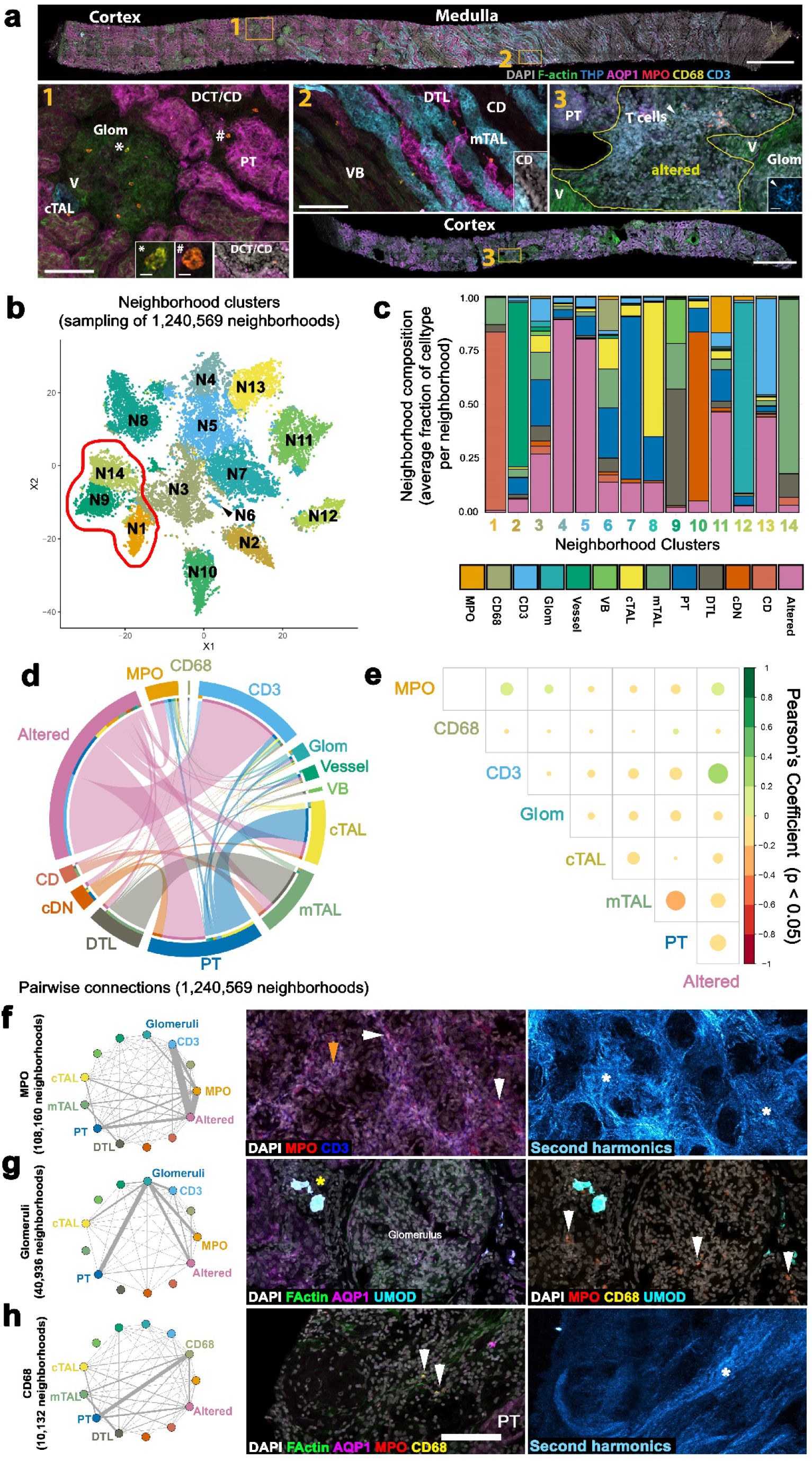
Defining cellular niches in renal disease from 3D fluorescence imaging. **a.** Maximum intensity projections of representative biopsies (cortex or medulla) showing classification label examples (insets **1-3).** These include: vessels (V, **1** and **3**) glomeruli (Glom, **1**), proximal tubules (PT, **1**), descending thin limb (DTL), medullary thick ascending limb (mTAL, **2**), vascular bundle (VB, **2**), cortical TAL (c-TAL, **1**), distal convoluted tubule, connecting tubules and collecting ducts (DCT/CNT/CD or cDN, **1**), medullary CD (CD, **2**) and areas of altered morphology or injury (altered, **3**). Examples of MPO+ and CD68+ are indicated in **1**. Scale bars are 1 mm in biopsy images, 100 um in **1** and **2** and 5 um in insets. **b**. Community based clustering on cell composition for ∼20,000 randomly chosen neighborhoods (15 biopsies or individuals). The red outline indicates neighborhoods including the medulla. **c**. Average cellular composition of the neighborhoods identified in (**b**). **d**. Pairwise analysis of cells within 1.2 million neighborhoods (15 biopsies or individuals), colors as indicated in (**c**). **e**. Pearson’s Coefficients for select interactions, the color indicates both the value and direction of the correlation. **f-g**. Neighborhoods with at least one cell for the labels indicated (MPO, Glomeruli and CD68) were subsetted and neighborhood graphs generated to indicate the pairwise interaction between cell labels. At right: maximum Z-projections of 3D confocal fluorescence images with white arrow indicating MPO+ cells (**f** and **g**) or CD68+ cells (**h**), orange arrows indicating CD3+ cells and asterisks highlighting fibrosis (white) or areas of altered morphology/injury (yellow). Scale bar = 100 um.

### Reference and Altered States

We now provide a higher level of complexity for all cell types along the depth of a kidney lobe from the cortex to the papillary tip (**Fig. 2a**), identifying 53 canonical human kidney cell types with associated biomarkers (**Supplementary Tables 8-9**). This includes a higher granularity for the loop of Henle, distal convoluted tubule and collecting duct segments, now resolving: three descending thin limb cell types (DTL1, 2, 3); different subpopulations of medullary thick ascending limb cells (M-TAL); two types of distal convoluted tubule cells (DCT1, 2); intercalated and principal cells of the connecting tubules (CNT-IC and CNT-PC); cortical, outer medullary and inner medullary collecting duct subpopulations (CCD, OMCD, IMCD); and papillary tip epithelial cells abutting the calyx (PapE). We further provide molecular profiles for several rare cell types important in homeostasis, including: juxtaglomerular renin-producing granular cells (REN); macula densa (MD); and a novel cell population enriched in schwann/neuronal (SCI/NEU) genes *NRXN1*, *PLP1* and *S100B* (**Supplementary Table 9**). We were further able to stratify: major endothelial cell types, including endothelial cells of the lymphatics (EC-LYM) and vasa recta (EC-AVR, EC-DVR); major stromal cell types including distinct fibroblast populations oriented along the cortico-medullary axis; and 12 immune cell types from lymphoid and myeloid lineages.

Through harmonized SNARE2 dual-omic annotations, we characterized the epigenetic landscape distinguishing the major kidney cell types found in snCv3 data in the cortex and medulla (**Extended Data Fig. 5**). Using paired AC data from the same nuclei annotated using RNA expression profiles, we identified open chromatin regions and candidate cis-regulatory elements for cell type marker genes, as well as associated transcription factor (TF) binding motif enrichments (**Extended Data Fig. 5a-b**, **Supplementary Tables 10-11**). We further identified accessibility of TF binding sites (TFBS), indicative of potential activity of expressed TFs, across most of the cell types identified by snCv3 (**Extended Data Fig. 5c, Supplementary Table 12)**. These include HNF4A in proximal tubule (PT), ESRRB in the TAL, GATA3 in the collecting tubules, FOXI1 in IC cells, SOX17 in ECs and MEF2D in VSMC/P.

**Figure 5.**
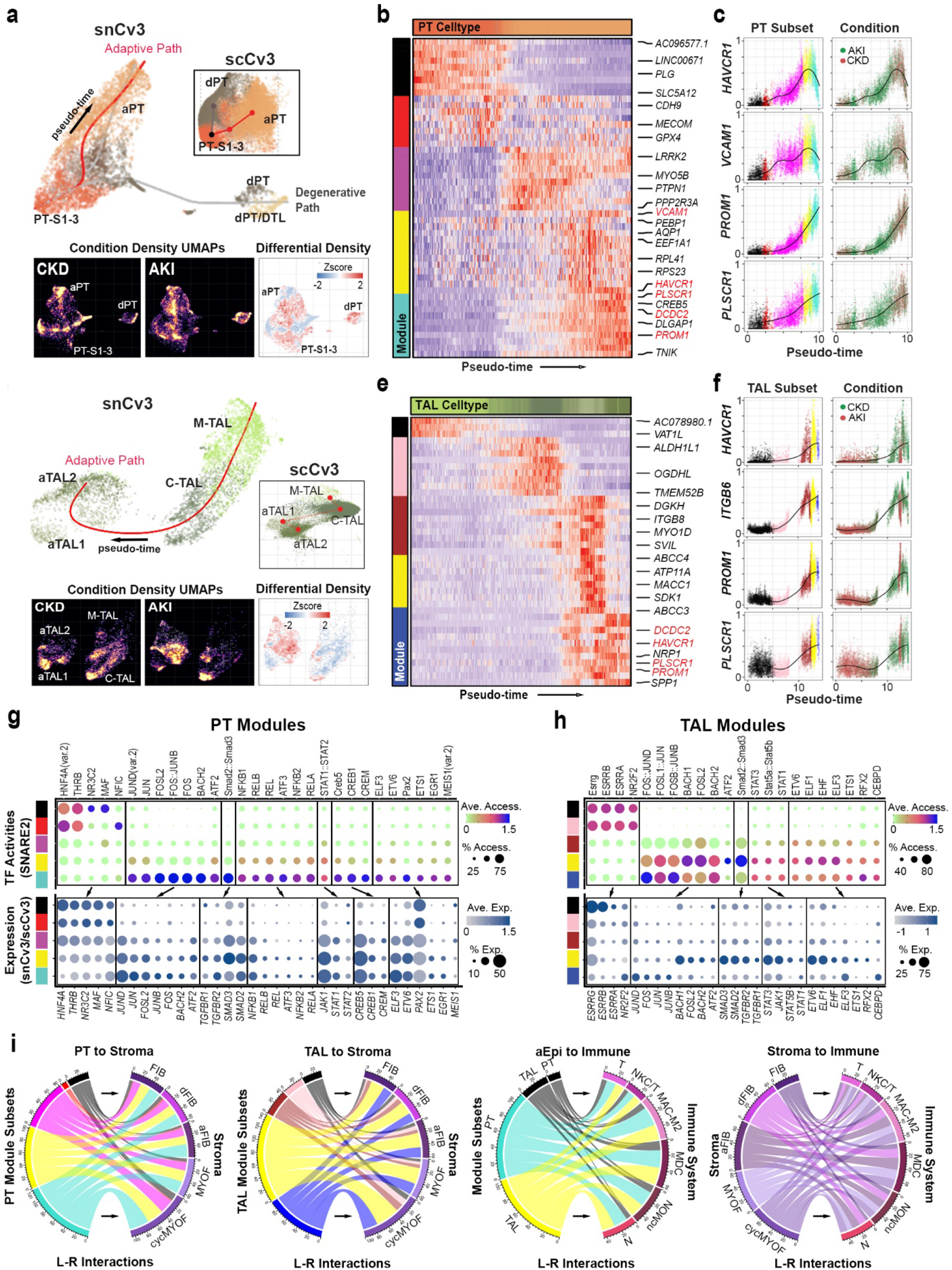
Expression and regulatory signatures of adaptive epithelial cells. **a.** Top: Trajectory of PT cells for snCv3 and scCv3 datasets. Bottom: PT embeddings colored based on cell density. The right panel shows the cell density difference between AKI and CKD. **b.** Heatmap of smoothed gene expression profiles along the inferred pseudo-time for PT cells. Color blocks on the left showing different modules identified based on the gene expression profiles. **c.** Left panels: changes of smoothed gene expression as a function of inferred pseudo- time colored based on the cells associated with their correspondent modules. Right panels: changes of smoothed gene expression as a function of inferred pseudo-time colored based on disease conditions. **d.** Trajectory of TAL cells for snCv3 and scCv3 datasets. Bottom: TAL embeddings colored based on cell density. The right panel shows the cell density difference between AKI and CKD. **e.** Heatmap of smoothed gene expression profiles along the inferred pseudo-time for TAL cells. Color blocks on the left showing different modules identified based on the gene expression profiles. **f.** Left panels: changes of smoothed gene expression for representative genes as a function of inferred pseudotime colored based on the cells associated with their correspondent modules. Right panels: changes of smoothed gene expression as a function of inferred pseudotime colored based on disease conditions. **g.** Top panel: dot plot of SNARE2 average accessibilities (chromVAR) and proportion accessible for TFBSs showing differential activity in aPT modules. Bottom panel: dot plot of averaged gene expression values (log scale) and proportion expressed for integrated snCv3/scCv3 modules. **h.** Dot plots as in (**g**) for aTAL modules. **i.** Circos plots showing number of secreted (non-integrin) ligand-receptor interactions between different cell populations. Arrows indicate direction of the interaction.

To spatially localize cell types within the tissue, snCv3 subclasses were used to predict the corresponding identities in Slide-seq and Visium transcriptomic data at different resolution scales (10µm and 55µm beads, respectively) (**Fig. 2c-i**, **Extended Data Fig. 6-7**, **Supplementary Table 2, Methods**). This allowed for recapitulated renal corpuscle, tubular, vascular, and interstitial cell types having proportions, marker profiles, and spatial organizations consistent with expected or observed (Visium) histopathology (**Extended Data Fig. 6-7**). Proximity network analysis based on the cell type composition of adjacent Slide-seq beads across 9 tissue pucks delineated cellular neighborhoods (**Fig. 2d**), including the renal corpuscle (RC) composition of podocytes (POD), glomerular capillaries (EC-GC), mesangial cells (MC), and parietal epithelial cells (PEC). These localized adjacent to the juxtaglomerular apparatus cells, REN and MD, and endothelial cells of the afferent/efferent arterioles (EC-AEA) leading into and out of the RC (**Fig. 2e-f**). This neighborhood analysis further identified a distinct vascular smooth muscle cell (VSMC) population juxtaposing or flanking the AEA (**Fig. 2g**).

**Figure 6.**
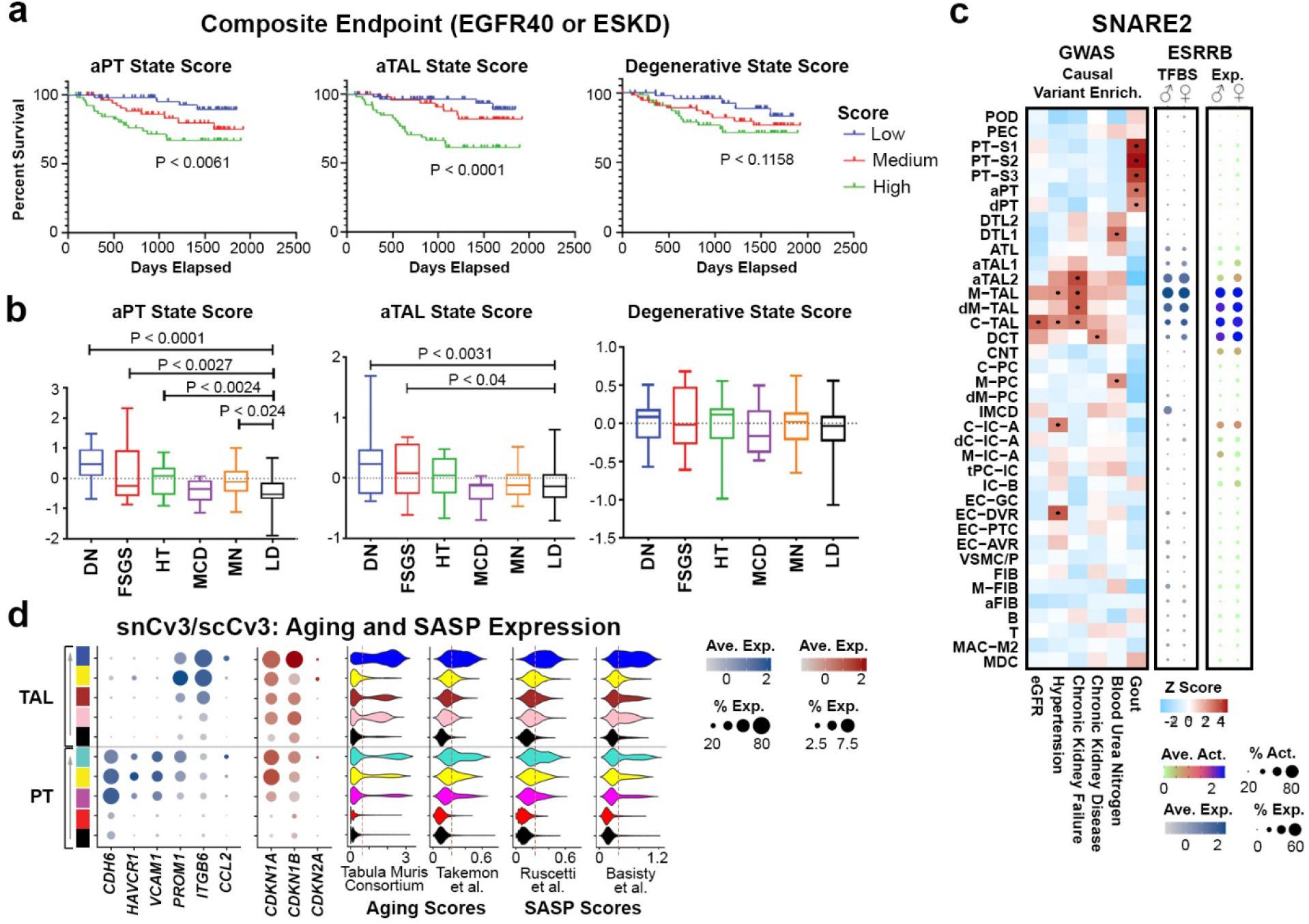
Adaptive signatures are associated with poor clinical outcome. **a.** Unadjusted Kaplan Meier curves by cell state scores for composite of end stage renal disease (ESRD) or for 40% drop in estimated glomerular filtration rate (eGFR) from time of biopsy in Neptune adult patient cohort (199 patients). Patients that reached the endpoint between screening and biopsy were excluded. The P values of log-rank tests for trend are shown. **b.** Boxplot of aPT, aTAL and degenerative state scores by kidney disease groups in the ERCB cohort (111 patients). Disease groups include diabetic nephropathy (DN), focal segmental glomerulosclerosis (FSGS), hypertensive nephropathy (HT), minimal change disease (MCD) and membranous nephropathy (MN). Boxes extend from the 25th to the 75th percentile for each group’s distribution and horizontal lines denote median values. Significant P values from unpaired t-tests between disease groups and living donors (LD) are shown. **c.** Heatmap of causal variants (z-scores) peak enrichments. Dots represent Z-scores > 2 (or P value < 0.05). Dotplots show averaged ESRRB binding site accessibility or gene expression (log values) and percent accessible or expressed. **d.** Dot plots of averaged gene expression values (log scale) and proportion expressed for integrated snCv3/scCv3 modules. Violin plots show gene expression scores for gene sets associated with aging or SASP (Methods).

Consistent with these annotations, we see the appropriate localization of associated cell type markers *REN* (REN), *NOS1* (MD), *NPHS2* (POD) and *MYH11* (VSMC), *SLC5A12* (PT-S1), *EMCN* (EC-GC) (**Fig. 2f-h**). In addition to the RC, we confirmed spatial resolution of subpopulations between the cortex and medulla, with the transition of C-TAL to M-TAL, both expressing *SLC12A1*, within the medullary rays (**Fig. 2i**). Therefore, the unique strengths of each spatial technology has enabled cross validation for our omic-defined cell type annotations. This permitted spatial localization of these cell types into functional tissue units, and more stratified annotations for distinct VSMC cell populations.

In addition to healthy states, a critical and novel element of this reference atlas is the characterization of cellular states associated with perturbations or injury. We carefully defined these altered states based on prior studies and gene expression profiles for clusters showing known features of injury (**Supplementary Table 13, Methods**). From this we established multiple putative states from cycling, transitioning, adaptive (or maladaptive) repair, to the degenerative (degen) states that may ultimately progress to necrosis or apoptosis. Applying these definitions, we identified altered states within snCv3 data for cell types found along the nephron, as well as within the stroma and vasculature (**Fig. 2, Supplementary Table 4**). These were contributed at different proportions from both reference and disease tissues and found to exist across technologies (**Extended Data Fig. 1**, **3**, **4**).

Clusters associated with the putative adaptive or maladaptive repair states were predominantly found within the PT and TAL subclasses, which may be due to the higher abundance of these tubules. Adaptive PT (aPT) clusters showed correlation with maladaptive states in rodents (**Extended Data Fig. 2e**), with characteristic expression of *VCAM1*, *DCDC2* and *HAVCR1* (**Extended Data Fig. 8a, Supplementary Table 14**)^3, 19^. Interestingly, we also identified a similar, as yet uncharacterized, state within the TAL, marked by *PROM1* (CD133) and *DCDC2* (**Extended Data Fig. 8a**). These are consistent with CD133+ PAX2+ lineage-restricted progenitors known to exist in the proximal and distal tubules of the adult kidney^20, 21^. Both of these adaptive epithelial (aEpi) cell types showed expression profiles associated with epithelial differentiation, morphogenesis and EMT, while also exhibiting a marked down-regulation of transporters critical to their normal function (**Extended Data Fig. 8b-c**). Furthermore, both aEpi cell types shared common signaling pathways and TF activities associated with injury related signaling, including mitogen-activated protein kinases (MAPKs) FOS/JUN, TGF-β and JAK/STAT^22^ (**Extended Data Fig. 8d**). This suggests a common aEpi state, sharing molecular signatures associated with injury and repair, that occurs in higher abundance within the PT and cortical TAL. Further, we find heterogeneity in aEpi clusters, with different developmental and differentiation pathways (aPT: SOX4, SOX6 and SOX13; aTAL: PAX2, TCF12 and PKNOX1) and distinct FOS/JUN and REL enriched clusters that may show distinct contributions to either successful or failed repair. We also identified separate adaptive states within the stroma (aStr) that are consistent with cell types contributing to wound healing and fibrosis following tissue injury (**Extended Data Fig. 2g**)^23^. These include myofibroblasts (MyoF), cycling MyoF (cycMyoF) and a population of adaptive fibroblasts (aFIB) representing potential MyoF progenitors^23^. We find increased expression of genes encoding periostin (*POSTN)*, fibroblast activation protein alpha (*FAP*), smooth muscle actin *(ACTA2)* and collagens, characteristic of these altered states (**Extended Data Fig. 9a**).

To assess altered state severity at the cellular level, we developed a scoring system using a strategy previously employed for single-cell ECM expression (**Extended Data Fig. 9**)^23^ using conserved genes upregulated in each of the altered states (degen, aPT, aTAL, aStr and cycling) across conditions (reference, AKI, CKD) (**Supplementary Tables 15-18**). Consistently, the state of cell clusters or subclasses within snCv3 and scCv3 could be predicted by their aggregate score values (**Extended Data Fig. 9b-e**). For example, aStr high scoring cell populations also showed high matrisome scores that is in line with their predicted role in ECM deposition. We also found elevated cycling state scores within AKI tissues compared to CKD (**Extended Data Fig. 9g**). This, and the potential enrichment of aEpi scores in AKI for a number of distal tubules, implies a higher level of repair or remodeling may be underway following acute injury events compared to ongoing chronic injury.

In addition to adaptive state signatures, we find common expression signatures that are shared across degenerative states coinciding with elevated expression of the known injury markers *SPP1, CST3*, *CLU* and *IGFBP7*^24^ (**Extended Data Fig. 9d-e**). Consistent with this, SNARE2 AC data identified common TFBS activities that may play a role in kidney cell degeneration, and that were associated with FOS/JUN signalling (**Extended Data Fig. 9f**). Therefore, common expression signatures associated with altered states permit single-cell/nucleus scoring, allowing both cellular level classification and possible insight into pathogenetic mechanisms of disease. Altered state scoring also provides a means for tagging injury populations in reference tissues arising from sample acquisition or normal aging, allowing for a cleaner representation of a healthy tissue reference atlas (**Extended Data Fig. 10**).

For spatial localization of injury, altered states were predicted along with reference states in both Slide-seq (aEpi, aStr, cycling) and Visium (aEpi, aStr, cycling, transitioning and degenerative) data. From Slide-seq, we identified areas of potential fibrosis around the AEAs that were enriched for aStr (aFIB, MyoF) and immune cell types, and which showed elevated *COL1A1* expression (**Fig. 3a-e**). We also identified an adjacent aTAL population with downregulated *EGF* expression, known to occur upon TAL injury^25^, and an upregulation of the aTAL marker *ITGB6* (**Fig. 3c, Supplementary Table 16**). For more detailed coverage of altered states, we used Visium on diseased tissues (**Figure 3f-k**), where there was an expected enrichment for adaptive states in CKD compared to reference tissues (**Extended Data Fig. 7b**). Furthermore, this technology permitted direct linkage of molecular profiles to histological areas of injury. Using this strategy, we interrogated an area of chronic fibrosis within a cortical CKD specimen (**Fig. 3f-g, Extended Data Fig 7e-f**). We found significant fibrosis that was associated with cell-type signatures arising from the stromal (FIB), aStr (aFIB), and immune cell clusters, especially monocyte derived cells (**Fig. 3g**, **Extended Data Fig. 7e-f**). There was also evident degeneration of FIB with increased expression of *B2M* and *VIM* (**Fig. 3g**). This region was surrounded by dilated and atrophic tubules that showed an aPT signature, including upregulation of *CDH6*^19^ (**Fig. 3g**, **Supplementary Table 16**). We also identified an area of PT- S1, showing degenerative and adaptive signatures, with *CDH6* expression adjacent to an area of MyoF accumulation and immune cell infiltration (**Fig. 3h-j**, **Extended Data Fig. 7g-h**).

In addition to cortical cell types, we found evidence for medullary injury of the collecting duct (**Fig. 3j-k**). Here we identified an arc of injured tubules, most with intraluminal cellular cast formation, cell sloughing, and loss of nuclei. This region was associated with degenerative CD cells, including dM-PC and transitioning principal and intercalated cells (tPC-IC) (**Fig. 3k**). Consistently, the degenerative marker *DEFB1* was locally up-regulated in this region where it may contribute to fibrosis by recruiting immune cells^26^. We also found distinct spatial localization of medullary vascular (EC-DVR, VSMC/P) and stromal (M-FIB) cell types adjacent to the region of injury (**Fig. 3j**). Therefore these results support co-mapping of reference and altered cell types identified from omic technologies, with specific states localized to histologic areas of injury in the appropriate cortical or medullary region of the kidney.

### Spatially Mapped Injury Neighborhoods

To uncover *in situ* cellular niches and injured microenvironments across kidney disease we probed the growing KPMP cohort of 3D imaging data of kidney biopsies (**Extended Data Fig. 11a, Supplementary Tables 2-3**). This included 3D fluorescence and second harmonic (fibrosis) image generation for specimens from both AKI and CKD patients (15 individuals, several interrogated by multiple technologies) and sampling of cortical and/or medullary renal tissue^27^. We used 3D-tissue cytometry to identify the composition of cellular niches associated with areas of altered or injured morphology. Cellular niches were defined for every cell (1,540,563 total over 15 individuals) by neighborhood analysis (cells within 25 µm) based on the 14 classes that covered the majority of renal cortical structures (**Fig 4a**, **Extended Data Fig. 11b**, Methods). From over 1.2 million total neighborhoods, we identified 14 unique groupings through community detection that included expected niches of cortical or medullary epithelium (N7 and N8 vs N14, N9 and N1 respectively, **Fig. 4b-c**). The TAL and PT epithelium neighborhoods (N7 and N8), as compared to other tubular epithelium and renal structures, had distinct neighborhoods enriched with areas of injury (**Fig. 4c** and **Extended Data Fig. 11c**). Furthermore, areas of injury were associated with infiltrating leukocytes including neighborhoods of CD68+, MPO+ and CD3+ cells (N6, N11 and N13 respectively). Uniquely, CD3+ cells were detected in a subset of neighborhoods almost exclusively with areas of tissue damage including presumptive epithelial degeneration (loss of markers and simplification) and fibrosis (N13, **Fig. 4c, a3 and Extended Data Fig. 11e**). In contrast, the myeloid cells were found in more cellular diverse niches including two neighborhoods with either cortical or medullary epithelium (N6 and N11, **Fig. 4c**). The leukocyte diversity was unique in these neighborhoods, as MPO+ and CD3+ cells were overlapping in neighborhoods (N11), whereas CD3+ cells were conspicuously low in neighborhoods with CD68+ cells (N6). Pairwise associations within neighborhoods identified a positive correlation between CD3+ and MPO+ but not CD68+ cells (**Fig. 4d-e**). Performing similar pairwise analyses for subsets of neighborhoods further identified positive correlations between leukocytes and specific renal structures, including CD68+ cells with PT epithelium and MPO+ cells with glomeruli (**Fig. 4e-h**). Overall, we found that altered states associated with renal injury in disease were enriched in PT and TAL neighborhoods, and showed predominantly CD3+ immune cell activity (**Fig. 4c**, **Extended Data Fig. 11c,e**). Thus, 3D imaging and tissue cytometry analysis of 1.2 million neighborhoods demonstrated distinct immune-active cellular niches and their association with discrete regions of healthy and injured tubules.

### Adaptive or Maladaptive Repair States

To obtain a deeper understanding of the genetic networks underlying the progression and potential pathology of altered PT and TAL, we performed trajectory inference on snCv3/SNARE2 and scCv3 subpopulations (**Fig. 5a-f**). While most degen states appeared too disconnected, both segment trajectories for the adaptive progression did show a transition from gene expression modules associated with normal function (black/red - PT; black/pink - TAL) to those associated with differentiation (magenta/yellow/turquoise - PT; brown/yellow/blue - TAL, **Supplementary Tables 19-20**). A majority of the expression gains were conserved across platforms (snCv3/SNARE2 and scCv3) and were found to occur towards the end of each trajectory (**Extended Data Fig. 12a-g**). These were associated with progenitor states that coincided with both maximal *PROM1* (CD133) expression (**Fig. 5 c, f**) and overlap with genes associated with failed repair in mouse AKI^3^ (turquoise module - PT, **Extended Data Fig. 12c**). There was also a concomitant increase in *HAVCR1* (KIM1) that was higher in PT, yet appeared elevated in AKI over CKD for TAL samples (**Fig. 5 c, f**). This suggests that this state, while potentially arising from acute injury, may persist in chronic disease.

Expression signatures across the trajectories revealed an enhancement in growth factor signaling with known roles in promoting tubulogenesis, maladaptive repair, fibrosis and inflammation. This includes Wnt (*DCDC2*, *PRICKLE*), Notch, TGF-β (*ITGB6*), EGF (*PLSCR1*) and Rho/Rac signalling pathways (**Fig. 5b,e**, **Extended Data Fig. 12d**, **Supplementary Tables 19-21**)^28–36^. Furthermore, we identify progressive activation of the MAPK (FOS/JUN), TGF-β and JAK/STAT pathways across both nephron segments, as predicted from TF activities associated with gene modules (**Extended Data Fig. 12i,k, Supplementary Table 22**) and TF motif accessibilities across adaptive trajectories (**Fig. 5 g-h, Supplementary Table 23**). Consistently, proximal tubule cells that showed expression of PROM1 were also found subjacent to phosphorylated JUN (p-JUN) likely suggesting close association of maladaptive and reparative cells (**Extended Data Fig. 12l-q**). As shown in prior studies, we identified progressively active REL/NF-KB signaling along the aPT trajectory^14^, that was also predicted based on expression modules in the aTAL trajectory (**Extended Data Fig. 12k**). We also found increased cAMP signaling (Creb TFs in aPT) capable of promoting dedifferentiatiation^37^ and increased ELF3 activities potentially required for MET^38^, both indicating that adaptive states may be poised for re-epithelialization. Therefore, we find adaptive epithelial trajectories sharing common molecular profiles that progressively upregulate cytokine signaling involved in tubule regeneration, while also providing molecular links to pathways associated with fibrosis, inflammation and end-stage kidney disease.

Given the upregulation of fibrotic cytokine signaling along adaptive trajectories, these regenerating cells may represent maladaptive states if they accumulate or fail to complete tubulogenesis. Therefore, we investigated the contribution of these states to cell-cell secreted ligand-receptor interactions within a fibrotic niche (**Supplementary Tables 24-26**). From imaging assays, this niche may comprise aEpi cells adjacent to normal and altered arteriole cells and fibroblasts, and immune cells that include T cells or macrophages depending on the level of tubular degeneration (**Figures 3-4**). Using snCv3 and scCv3 data sets associated with trajectory modules, we identified both late aPT and aTAL states as having a higher number of interactions with the stroma (**Fig. 5i**). This was associated with secreted growth factors of the FGF, BMP, WNT, EGF, IGF and TGF-β families (**Extended Data Fig. 13a-b**). Furthermore, late modules and aStr cell types showed a higher number of ligand-receptor interactions with immune cells (**Fig. 5i**, **Extended Data Fig. 13c-d**). This indicates adaptive tubule states may recruit immune cells both primarily and secondarily through their recruitment of the activated fibroblasts and myofibroblasts. This is consistent with the activation of Rel/NF-kB and CEBPD transcription factors, having known roles in promoting inflammation ^39, 40^, in the aEpi populations (**Fig. 5g-h**). We also found expression of the PVR Cell Adhesion Molecule (CD155) gene in late aTAL modules that may mediate its interactions with natural killer (NK) cells, or provide a mechanism to escape immune surveillance through PVR association with TIGIT (**Extended Data Fig. 12e**)^41, 42^. The upregulation of PVR in aTAL and not aPT might contribute to the fewer observed T or NKC/T cell associations with C-TAL compared to PT neighborhoods (**Fig. 4e-f**).

We also find additional evidence for the activation of EGF pathway signaling within the adaptive epithelial trajectories, which in itself may lead to activation of TGF-β signalling and create a niche capable of promoting fibrosis^36^. Consistently, EGF ligands NRG1 and NRG3 both become expressed in aEpi states for a possible role in MAC-M2 recruitment (**Extended Data Fig. 12c,e**). Furthermore, expression of EGF receptors ERBB2, ERBB4 (aPT/aTAL) and ERBB3 (aPT) may poise these cells for contribution to autocrine/paracrine signalling within the adaptive tubules (**Extended Data Fig. 12e**). Since MAPK pathways can mediate ErbB receptor signaling, it remains possible that the increased activity of FOS/JUN could in fact be associated with EGF pathway functions promoting regeneration (**Fig.5g-h**). Therefore, we identify expression and regulatory signatures associated with a common reparative state in proximal and distal tubules. However, this may represent a maladaptive state that produces and receives a number of cytokine signals that promotes both fibrosis and inflammation. In support of this, we find PROM1 expression along either trajectory to be elevated within CKD compared to AKI cases (**Fig. 5c,f**). We also find distinct expression profiles exist within different tubular segments that may modulate how these cells interact with their fibrotic niches or contribute to disease progression.

### Adaptive but not degenerative state scores associate with progressive decline in kidney function

To identify whether aEpi cell states contribute to chronic kidney disease, we identified gene signatures for altered states that were conserved across technologies (snCv3 and scCv3) (**Supplementary Table 27**) and that were associated with disease severity (**Extended Data Fig. 14a-d**). These signatures were assessed for their association with disease progression within the Nephrotic Syndrome Study Network (NEPTUNE) cohort of 199 patients^43^. Composite gene expression scores were computed on the tubulointerstitial compartment for degenerative and adaptive cell states and used for Kaplan-Meir (K-M) analyses. In an unadjusted survival model, high adaptive, but not degenerative, state scores were significantly associated with composite endpoint (40% loss of eGFR or ESKD), with aTAL and aStr showing the most significant associations (p value < 0.0001) (**Fig. 6a,Extended Data Fig. 14e**). This indicated that aEpi processes may represent maladaptation and, like fibrosis-promoting aStr states, associate with disease progression. Alternatively, degenerative states progressing to necrosis or apoptosis may not accumulate over time. Interestingly, high adaptive state scores from a common set of aPT-aTAL genes were also found to have a significant association with faster end point (p value < 0.0015), indicating a common, adaptive epithelial state that may accumulate or persist and ultimately contribute to eventual organ failure due to maladaptive repair.

Additional analysis of transcriptomic data from 111 kidney disease patients in the European Renal cDNA Bank (ERCB) cohort^44^, found scores for all adaptive, but not degenerative, states were significantly higher in the diabetic nephropathy (DN) patients compared to that of living donors (LD) (**Fig. 6b, Extended Data Fig. 14f**). The high association with ESKD and DN scores were found for each adaptive tubule type, demonstrating critical roles for effective repair mechanisms not only in the PT, but also in the TAL.Therefore, TAL functionality, which may include its known GFR-regulatory role through tubuloglomerular feedback, may represent a major contributing factor to progressive kidney failure. Consistent with this, causal variants for eGFR and chronic kidney failure were found to be enriched within TAL regulatory regions that also were enriched for Estrogen Related Receptor (ESSR) TF motifs (**Fig. 6c, Supplementary Table 28**). ESRR TFs (especially ESRRB), key players in TAL ion transporter expression^45^, are central regulators of the TAL expression network (**Extended Data Fig. 14g)** and become inactivated in adaptive states (**Fig. 4h**). Therefore, we demonstrate both a potential maladaptive role for the aEpi states and a potential central role for the TAL segment in maintaining the health and homeostasis of the human kidney. This is consistent with the finding that the top renal genes showing decline in a mouse aging cell atlas were associated with the TAL^46^.

Our findings implicate an accumulation of maladaptive epithelia during disease progression that may be consistent with chronically senescent cells^4^. This is supported by both increased expression of aging related genes and an apparent senescence-associated secretory phenotype (SASP) for these cells (**Fig. 5, Fig. 6d, Extended Data Fig. 14h)**. As such, we detected *CDKN1A* (p21^cip1^), *CDKN1B* (p27^kip1^), *CDKN2A* (p16^ink4a^) and *CCL2* expression in late aPT and aTAL states (**Fig. 6d**). Furthermore, expression signatures for reparative processes in aEpi states were downregulated in the CKD (n = 28) over AKI (n = 22) cases used in this study (snCv3/scCv3), while G1/S checkpoint regulatory factors were upregulated (**Supplementary Table 30**). This is consistent with repair processes that may persist after injury^19^, but that may subsequently transition to senescent pro-fibrotic states during disease progression.

## Discussion

We present a comprehensive spatially resolved cell atlas to define genes and pathways across the corticomedullary axis of the kidney, including signalling between tubules, stroma and immune cells that underlie normal and pathological cell neighborhoods. Through careful definition of injury states, we identify putative adaptive or maladaptive repair signatures within the epithelial segments that may reflect a failure to complete differentiation and tubulogenesis. This enabled us to resolve and greatly expand upon existing healthy reference and altered state cell identities. Spatial analyses prioritized relevant cell-cell interaction niches associated with altered injury states and permitted reconstruction of the fibrotic niche. From this we find that expression signatures for the progression of adaptive states within the proximal and distal tubules are associated with elevated cytokine production, increased interactions with the fibrotic and inflammatory cell types and ultimately the progression to end stage kidney disease. These adaptive state signatures were highly associated with tubule regeneration and differentiation, indicating that the potential failure of these cells to complete tubulogenesis might ultimately lead to a progressive decline in kidney function. This may arise from an incompatible mileu associated with the high level of cytokine signalling found within the fibrotic niche. In turn, the high cytokine producing nature of these cells may further contribute to kidney disease through promotion of fibrosis. We identified specific modules in aEpi states enriched in senescence associated genes suggesting likely perturbation of cell cycle progression that will require deeper evaluation. Since several adaptive markers were overlapped across tubular regions, physiological or pathological stresses may initiate activation of common signaling events that could be subject to the same therapeutic strategies.

In this study, we have leveraged multiple technologies, samples, sites and health conditions, representing efforts between the HuBMAP, KPMP and HCA consortia, to define cell types and states underlying health and disease. This atlas will serve as a key resource for studies into: normal physiology and sex differences; pathways associated with transitions from healthy and injury states; clinical outcomes; disease pathogenesis; and targeted interventions.

## Methods

### Statistics and Reproducibility

For spatial transcriptomics, 3D imaging and immunofluorescence staining experiments, each staining was repeated on at least 2 separate individuals or separate regions. For SLIDE-seq where only one individual was available, the assay was performed on 9 adjacent tissue sections. For immunofluorescence validation studies, commercially available antibodies were used; the immunostaining included tissue from patients not contributing to omics data. Similarly, orthogonal validation of omics annotations and spatial localization in Visium studies also included more than four samples each from reference and disease biopsies that were not used to generate single cell gene expression data to further increase the reproducibility and rigor. Further, several technologies were performed on samples from the same patient and in some cases the same tissue block was used to generate multimodal data.

### Ethical Compliance

We have complied with all ethical regulations related to this study. Human samples (**Supplementary Table 1**) collected as part of the Kidney Precision Medicine Project (KPMP) consortium (KPMP.org) were approved as exempted by the University of Washington Institutional Review Board. Samples as part of the Human Biomolecular Atlas Program (HuBMAP) consortium were collected by the Kidney Translational Research Center (KTRC) under a protocol approved by the Washington University Institutional Review Board (IRB *#*201102312). Informed consent was obtained for the use of data and samples for all participants at Washington University, including living patients undergoing partial or total nephrectomy or from discarded deceased kidney donors. For Visium Spatial Gene Expression, reference nephrectomies and diabetic kidney biopsy specimens were obtained from the KPMP or the Biopsy Biobank Cohort of Indiana (BBCI)^47^ as approved by the Indiana University Institutional Review Board (IRB # 1906572234). Living donor biopsies as part of the Human Cell Atlas (HCA) were obtained under the Human Kidney Transplant Transcriptomic Atlas (HKTTA) under IRB HUM00150968. Deidentified leftover frozenCOVID-19 AKI kidney biopsies were obtained from the Johns Hopkins University under IRB 00090103.

### Human Tissue Specimens

For single nucleus omic assays, tissues were processed according to the following protocol: dx.doi.org/10.17504/protocols.io.568g9hw. For nuclei preparation, ∼7 sections of 40 µm thickness were collected and stored in RNAlater solution (RNA assays) or kept on dry ice (AC assays) until processing or used fresh. To confirm tissue composition, 5 µm sections flanking these thick sections were obtained for histology and the relative amount of cortex or medulla composition including glomeruli was determined. For single cell omic assays, tissues used (15 CKD,12 AKI and 18 LD biopsy cores) were preserved using CryoStor® (Stemcell Technologies).

### RNA-Sequencing, QC and Clustering

#### Isolation of single nuclei

Nuclei were isolated from cryosectioned tissues according to the following protocol: dx.doi.org/10.17504/protocols.io.ufketkw with the exception that 4’,6-diamidino-2-phenylindole (DAPI) was excluded from the nuclear extraction buffer and only used to stain a subset of nuclei used for counting. Nuclei were used directly for omic assays.

#### Isolation of single cells

Single cells were isolated from frozen tissues according to the following protocol: dx.doi.org/10.17504/protocols.io.7dthi6n. The single cell suspension was immediately transferred to the University of Michigan Advanced Genomics Core facility for further processing.

#### 10X Chromium v3 (Cv3) RNA-sequencing

10X single nucleus RNA sequencing was performed according to dx.doi.org/10.17504/protocols.io.86khzcw, and the 10X single cell RNA sequencing according to dx.doi.org/10.17504/protocols.io.7dthi6n, both using the 10X Chromium Single-Cell 3’ Reagent Kit v3. Sample demultiplexing, barcode processing, and gene expression quantifications were performed with the 10X Cell Ranger v3 pipeline using the GRCh38 (hg38) reference genome. For single nucleus data, introns were also included in the expression estimates.

#### SNARE-Seq2 dual RNA and ATAC-sequencing

SNARE-Seq2^16^, as outlined (Nature Protocols, DOI:10.1038/s41596-021-00507-3), was performed according to the following protocol: dx.doi.org/10.17504/protocols.io.be5gjg3w. AC and RNA libraries were sequenced separately on the NovaSeq 6000 (Illumina) system using the 300 cycle and 200 cycle reagent kits, respectively.

#### SNARE-Seq2 Data Processing

Detailed step-by-step processing for SNARE-Seq2 data has been outlined (Nature Protocols, DOI:10.1038/s41596-021-00507-3). This has now been implemented as an automated data processing pipeline that is available at github.com/huqiwen0313/snarePip. The pipeline provides an automated framework for complex single-cell analysis including quality assessment, doublet removal, cell clustering and identification, robust peak generation and differential accessible region identification with flexible analysis modules and generating summary reports for both quality assessment and downstream analysis. The directed acyclic graph was used to incorporate the entire data processing steps for better error control and reproducibility. For RNA processing, this involved removal of AC contaminating reads using cutadapt (version 3.1) ^48^, dropEst (version 0.8.6) ^49^ to extract cell barcodes and STAR (v2.5.2b) ^50^ to align tagged reads to the genome (GRCh38). For AC data, this involved snaptools (version v1.2.3) ^51^ and minimap (version 2-2.20)^52^ for alignment to the genome (GRCh38).

#### Quality control of sequencing data

10X snRNA-seq (snCv3): Cell barcodes passing 10X Cell Ranger filters were used for downstream analyses. Mitochondrial transcripts (MT-*) were removed, doublets were identified using the DoubletDetection software (v2.4.0)^53^ and removed. All samples were combined across experiments and cell barcodes having greater than 200 and less than 7500 genes detected were kept for downstream analyses. To further remove low quality datasets, a gene UMI ratio filter (gene.vs.molecule.cell.filter) was applied using Pagoda2 (github.com/hms-dbmi/pagoda2). 10X scRNA-seq (scCv3): As a quality control step, a cutoff of < 50% mitochondrial reads per cell was applied. The ambient mRNA contamination was corrected using SoupX (v1.5.0)^54^.The mRNA content and number of genes for doublets are comparatively higher than for single cells. In order to reduce doublets or multiplets from the analysis, we used a cutoff of > 500 and < 5000 genes per cell.

SNARE-Seq2 RNA: Cell barcodes for each sample were retained with the following criteria: having DropEst cell score greater than 0.9; having greater than 200 UMI detected; having greater than 200 and less than 7500 genes detected. Doublets identified by both DoubletDetection (v3.0) and Scrublet (github.com/swolock/scrublet, version 0.2.2) were removed. To further remove low quality datasets, a gene UMI ratio filter (gene.vs.molecule.cell.filter) was applied using Pagoda2.

SNARE-Seq2 ATAC: Cell barcodes for each sample that had already passed quality filtering from RNA data were further retained with the following criteria: having tss enrichment greater than 0.15; having at least 1000 read fragments and at least 500 UMI; having fragments overlapping the promoter region ratio of greater than 0.15. Samples were only retained if they exhibited greater than 500 dual omic cells after quality filtering.

#### Clustering snCv3

Clustering analysis was performed using pagoda2, where counts were normalized to the total number per nucleus, batch variations were corrected by scaling expression of each gene to the dataset-wide average. After variance normalization, all 5526 significantly variant genes were used for principal component analysis. Clustering was performed at different k values (50, 100, 200, 500) based on the top 50 principal components, with cluster identities determined by the infomap community detection algorithm. The primary cluster resolution (k = 100) was chosen based on the extent of clustering observed. Principal components and cluster annotations were then imported into Seurat (version 4.0.0) and uniform manifold approximation and projection (UMAP) dimensional reduction was performed using the top 50 principal components identified using pagoda2. Subsequent analyses were then performed in Seurat. A cluster decision tree was implemented to determine whether a cluster should be merged, split further or labeled as an altered state. For this, differentially expressed genes between clusters were identified for each resolution using the FindAllMarkers function in Seurat (only.pos = TRUE, max.cells.per.ident = 1000, logfc.threshold = 0.25, min.pct = 0.25). Possible altered states were initially defined for clusters having one or more of the following features: low genes detected, high number of mitochondrial transcripts, high number of ER associated transcripts, upregulation of injury markers (*CST3, IGFBP7, CLU, FABP1, HAVCR1, TIMP2, LCN2*) or enrichment in AKI or CKD samples. Clusters (k = 100) that showed no distinct markers were assessed for altered state features, if present then these clusters were tagged as possible altered states, if absent then clusters were merged based on their cluster resolution at k = 200 or 500. If this merging would occur across major classes (epithelial, endothelial, immune, stromal) at higher k values, then these clusters were instead labeled as ambiguous or low quality (including possible multiplets). For k = 100 clusters (non-epithelial only) that did show distinct markers, their k = 50 subclusters were assessed for distinct marker genes, if present, then these clusters were split further. The remaining split and unsplit clusters were then assessed for altered state features. If present they were tagged as possible altered states, if absent they were assessed as the final cluster. Annotations of clusters were based on known positive and negative cell type markers^10, 11, 55–57^ (also see **Supplementary Table 5**), regional distribution of the clusters across the corticomedullary axis and altered state (including cell cycle) features. For separation of EC-DVR from EC-AEA, the combined population was independently clustered using pagoda2 and clusters associated with medullary sampling were annotated as EC-DVR. For separation of the REN cluster, stromal cells expressing *REN* were selected based on normalized expression values greater than 3.

#### Annotating snCv3 Clusters

To overcome the challenge of disparate nomenclature for kidney cell annotations, we leveraged a cross-consortium effort to use the extensive knowledge base from human and rodent single-cell gene expression data sets, as well as the domain expertise from pathologists, biologists, nephrologists and ontologists^10, 11, 19, 55–58^ (also see **Supplementary Table 4, 5** and the HuBMAP ASCT+B Reporter: hubmapconsortium.github.io/ccf-asct-reporter). This allowed the adoption of a standardized anatomical and cell type nomenclature for major and minor cell types and their subclasses (**Supplementary Table 4**), showing distinct and consistent expression profiles of known markers and absence of specific segment markers for some of the cell types (**Extended Data Fig. 2a, Supplementary Table 5**). The knowledge of the regions dissected and histological composition of snCv3 data further enabled stratification of distinct cortical and outer and inner medullary cell populations (**Fig. 2b, Extended Data Fig. 1**). The cell type identities and regional locations were confirmed through orthogonal validation using spatial technologies presented here and correlations with existing human or rodent stromal, immune, endothelial and epithelial data sets^3, 23, 55, 56, 58, 59^ (**Extended Data Fig. 2b-i)**.

### Integrating snCv3 and SNARE2 data sets

Integration of snCv3 and SNARE RNA data was performed using Seurat (v4.0.0) using snCv3 as reference. All counts were normalized using sctransform, anchors were identified between data sets based on the snCv3 pagoda2 principal components. SNARE2 data was then projected onto the snCv3 UMAP structure and snCv3 cell type labels were transferred to SNARE2 using the MapQuery function. Both data sets were then merged and umap embeddings recomputed using the snCv3 projected principal components. Integrated clusters were identified using pagoda2, with the k-nearest neighbor graph (k = 100) based on the integrated principal components and using the infomap community detection algorithm. The SNARE2 component of the integrated clusters was then annotated to the most overlapping, correlated and/or predicted snCv3 cluster label, with manual inspection of cell type markers used to confirm identities. Integrated clusters that overlapped different classes of cell types were labeled as ambiguous or low quality clusters.

### Integrating snCv3 and scCv3 data sets

Integration of snCv3 and scCv3 data was performed using Seurat (v4.0.0) using snCv3 as reference. All counts were normalized using sctransform, anchors were identified between data sets based on the snCv3 pagoda2 principal components. scCv3 data was then projected onto the snCv3 UMAP structure and snCv3 cell type labels were transferred to scCv3 using the MapQuery function. Both data sets were then merged and umap embeddings recomputed using the snCv3 projected principal components. Integrated clusters were identified using pagoda2, with the k-nearest neighbor graph (k = 100) based on the integrated principal components and using the infomap community detection algorithm. The scCv3 component of the integrated clusters was then annotated to the most overlapping or correlated snCv3 subclass, with manual inspection of cell type markers used to confirm identities. Cell types that could not be accurately resolved (PT-S1/PT-S2 and EC-AEA/EC-DVR) were kept merged. Integrated clusters that overlapped different classes of cell types or that were too ambiguous to annotate were considered low quality and were removed from the analysis.

### NSForest marker genes

To identify a minimal set of markers that can identify snCv3 clusters and subclasses (subclass.l3), or scCv3 integrated subclasses (subclass.l3), we used the Necessary and Sufficient Forest^60^ (NSForest v2, github.com/JCVenterInstitute/NSForest/releases/tag/v2.0) software using default settings.

### Correlation analyses

For correlation of RNA expression values between snCv3 and scCv3, or SNARE2, average scaled expression values were generated, pairwise correlations performed using variable genes identified from Pagoda2 analysis of snCv3 (top 5526 genes). For comparison with mouse single cell RNA-seq on healthy reference tissue^56^, raw counts were downloaded from the Gene Expression Omnibus (GEO, GSE129798). For comparison with mouse single cell RNA-seq from ischemia–reperfusion injury (IRI) tissue^3^, raw counts were downloaded from GEO (GSE139107). For human fibroblast and myofibroblast data^23^, raw counts were downloaded from Zenodo (10.5281/zenodo.4059315). For each data set, raw counts were processed using Seurat: counts for all cell barcodes were scaled by total UMI counts, multiplied by 10,000 and transformed to log space. For comparison with mouse single cell types of the distal nephron^58^, the precomputed Seurat object was downloaded from GEO (GSE150338). For mouse bulk distal segment data^58^, normalized counts were downloaded from GEO (GSE150338) and added to the “data” slot in a Seurat object. Immune cell reference data was obtained using the celldex package^61^ using the MonacoImmuneData()^59^ and ImmGenData()^61, 62^ functions and log counts imported into the “data” slot of Seurat. For correlation against these reference data sets, averaged scaled gene expression values for each cluster were calculated (Seurat) using an intersected set of variable genes identified for each data set (identified using Padoda2 for snCv3 and Seurat for reference data sets). For immune reference correlations, a list of immune-related genes downloaded from ImmPort (immport.org) was used instead of the variable genes. Only fine resolution immune labels having correlation greater than 0.2 were combined at the main label level for final correlation. Correlations were plotted using the corrplot package (github.com/taiyun/corrplot). Several of the immune annotations were further confirmed by manual comparison with recently reported data^13^.

### Computing single nucleus/cell-level expression scores

To identify markers associated with altered states (degenerative or degen; adaptive - epithelial or aEpi; adaptive - stromal or aStr; cycling), snCv3 and scCv3 data were independently used to identify differentially expressed genes between reference and corresponding altered states for each subclass level 1 (subclass.l1). To ensure general state-level markers, differentially expressed genes were identified using the “FindConservedMarkers” function (grouping.var = "condition.l1", min.pct = 0.25, max.cells.per.ident = 300) in Seurat. A minimal set of general degenerative conserved genes were identified as enriched (p value < 0.05) in the degenerative state of each condition.l1 (reference, AKI and CKD) and in at least 4 of the 11 (snCv3) or 9 (scCv3) subclass.l1 cell groupings. A minimal set of conserved aEpi genes were identified as enriched (p value < 0.05) in the adaptive state of each condition.l1 (reference, AKI and CKD) in both aPT and aTAL cell populations. This aEpi gene set was then further trimmed to include only those genes that were enriched within the adaptive epithelial population (aPT/aTAL) versus all others using the “FindMarkers” function and a minimum p value of 0.05 and average log2 fold change > 0.6. A minimal set of conserved aStr genes were identified as enriched (p value < 0.05) in the adaptive state of each condition.l1 (reference, AKI and CKD for snCv2; reference and AKI for scCv3) for stromal cells. To increase representation from MYOF in scCv3 showing a small number of these cells, MYOF-alone enriched genes (average log2 fold change >= 0.6; adjusted p value < 0.05) were included for the scCv3 gene set. The aStr gene sets were then further trimmed to include only those genes that were enriched within the adaptive stromal population (aFIB and MYOF) compared to all others using the “FindMarkers” function and a minimum p value of 0.05 and average log2 fold change > 0.6. A minimal set of cycling- associated genes were identified as those enriched (adjusted p value < 0.05 and average log2 fold change > 0.6) in the cycling state across all associated subclass.l1 cell groupings.

Scores for altered state, ECM and for gene sets associated with aging or SASP were computed for each cell from averaged normalized counts using only the genes showing a minimum correlation to the averaged whole gene set of 0.1^23^ (github.com/mahmoudibrahim/KidneyMap). Aging and SASP genes were obtained from the Tabula Muris Consortium (top 20 genes upregulated in aging kidney)^46^, Takemon et al. (genes from Table S3, group.age A^63^), Ruscetti et al.(SASP genes from Figure 2c)^64^ or Basisty et al.(from Table S1 sheet IR Epithelial SASP, having a positive AVE log2 ratio)^65^.

### Gene Set Enrichment Analyses (GSEA)

To compute gene set enrichments for aPT and aTAL, conserved genes differentially expressed in the adaptive over reference states were identified as indicated above. Gene set ontologies from the Molecular Signatures Database (MSigDB) were downloaded from gsea-msigdb.org and pathway enrichments computed using fgsea^66^ and gage^67^, keeping only GO that were significant (p < 0.05) for both. Redundant pathways were collapsed using the fgsea function “collapsePathways” and visualized using the ggplot.

### SNARE2 AC analyses

SNARE2 chromatin data was analysed using Signac^68^ (v1.1.1). Peak calling was performed using the “CallPeaks” function and MACS (v3.0.0a6, github.com/macs3-project/MACS) separately for clusters, subclass.l1 and subclass.l3 level annotations. Peak regions were then combined and used to generate a peak count matrix using the “FeatureMatrix” function, then used to create a new assay within the SNARE2 Seurat object using the “CreateChromatinAssay” function. Gene annotation of the peaks was performed using “GetGRangesFromEnsDb(ensdb = EnsDb.Hsapiens.v86)”. TSS enrichment, nucleosome signal and blacklist fractions were all computed using Signac. Jaspar motifs (JASPAR2020, all vertebrate) were used to generate a motif matrix and motif object that was added to the Seurat object using the “AddMotifs” function. For motif activity scores, chromVAR^69^ (v1.12.0, greenleaflab.github.io/chromVAR) was performed using the “RunChromVAR” function. The chromVAR deviation score matrix was then added to a separate assay slot of the Seurat object. For visualization of the chromatin data, UMAP embeddings were computed from cis-regulatory topics that were identified through Latent Dirichlet Allocation (LDA) using CisTopic^70^ (v0.3.0) (github.com/aertslab/cisTopic) and the “runCGSModels” function. Only regions accessible in 50 nuclei and nuclei having 200 of these accessible regions were used for cisTopic and downstream analyses. The umap coordinates for the remaining nuclei were added to the Seurat object. To ensure high quality AC profiles, only clusters having more than 50 nuclei were retained for downstream analyses (**Supplementary Table 7**).

### Differentially Accessible Site (DAR) analyses

Sites that were differentially accessible for a given cell grouping (subclass) were identified against a selection of background cells having best matched total peak counts, in order to best account for technical differences in the total accessibility for each cell. For this, the total peaks in each cell were used for estimation of the distribution of total peaks (depth distribution) for the cells belonging to the test cluster, and 10,000 background cells having a similar depth distribution as the test cluster were randomly selected. DARs were then identified as significantly enriched in the positive cells over selected background cells using the “CalcDiffAccess” function (github.com/yanwu2014/chromfunks), where p-values were calculated using a Fisher’s Exact Test on a hypergeometric distribution and adjusted p-values (or q-values) were calculated using the Benjamini & Hochberg (BH) method. For subclass level 2 DARs, VSM/P clusters were merged and the MD was combined with C-TAL prior to DAR calling. Subclasses having >100 DARs with q value < 0.01 were used for further analysis. Co-accessibility between all peak regions was computed using Cicero^71^ (v1.8.1). Sites were then linked to genes based on co-accessibility with promoter regions, occurring within 3000 base pairs of a gene’s transcriptional start site (TSS), using the “RegionGeneLinks” function (github.com/yanwu2014/chromfunks) and the ChIPSeeker package^72^. DARs associated with markers for each subclass (identified at the subclass.l2 level using snCv3, p value < 0.05) and showing q value < 0.01 and log fold change > 1 were selected for visualization. For this, DAR accessibility (peak counts) were averaged, scaled (trimming values to a minimum of 0 and a maximum of 5) and visualized using the ggHeat plotting function of the SWNE package^73^. Motif enrichment within cell type DARs were computed using the hypergeometric test (“FindMotifs” function) in Signac.

### Transcription factor analyses

To identify active TFs from SNARE2 AC data, differential activities (or deviation scores) of TFBS between different populations were assessed using the “Find[All]Markers” function through logistic regression and using the number of peak counts as a latent variable. Only TFs with expression detected within the corresponding cluster, subclass or state grouping were included. For PT and TAL clusters, TFBSs that were differentially active (p value < 0.05, average log2 fold change > 0.35) and associated with TFs with expression detected in at least 2.5% of nuclei (SNARE2) were identified between clusters. Common aPT/aTAL TFBS activities were identified from an intersection of those differentially active and expressed within adaptive PT and TAL clusters. For aPT and aTAL trajectory modules, TFBSs showing differential activity between modules (adjusted p value < 0.05, average log2 fold change > 0.35) and expression detected within at least 2.5% of nuclei/cells (snCv3/scCv3) were identified. For common degenerative state TFBS activities, differentially active TFBS were identified between reference and degenerative states for each level 1 subclass (**Supplementary Table 13**). Significant degenerative state TFBS activities (p value < 0.05, average log2 fold change > 0.35) in 3 or more subclass.l1 were trimmed to those showing expression detected in more than 20 percent of the degenerative state nuclei/cells for snCv3/scCv3.

### Ligand-receptor interaction analyses

Ligand-receptor analyses were performed using the CellPhoneDB python package (v2.1.7, github.com/Teichlab/cellphonedb) by running the statistical method on select subclasses or trajectory (aPT, aTAL) modules. Only interactions for secreted ligands that were not associated with integrins were visualized using ggplot. Ordering of the ligand-receptor interactions was based on hierarchical clustering (ward.D2 method) using the ggdendro (v0.1.20) package. Circos plots to summarize the number of interactions from one subclass subset to another were performed using the circlize package (github.com/jokergoo/circlize).

### Plots and figures

All UMAP, feature, dot, and violin plots for snCv3, scCv3 and SNARE2 data were generated using Seurat. Correlation plots were generated using the corrplot package. Genome coverage plots were performed using Signac. Plots for 3D cytometry and neighborhood analysis were generated in R with circois, ggplot2, and igraph.

### GWAS analyses

To link SNARE2 cell types to kidney GWAS traits and diseases, we first summed the binary peak accessibility profiles for all cells belonging to the same cell type to create a pseudobulk peak-by-subclass accessibility matrix. Pseudobulk analyses give more stable results, especially since SNARE2 accessibility data can be sparse. To ensure sufficient coverage, we used subclass level 2 groupings with the following modifications: VSM/P clusters were merged; MD was combined with C-TAL; subclasses having <100 DARs with q value < 0.01 were excluded. We used g-chromVAR^74^ (v0.3.2), an extension of chromVAR for GWAS data, to identify cell types with higher than expected accessibility of genomic regions overlapping GWAS-linked SNPs. Running g-chromVAR requires first identifying GWAS-linked SNPs that are more likely to be causal, a process known as fine-mapping. For the Chronic Kidney Failure GWAS traits, we used existing fine-mapped SNPs from the CausalDB database, using the posterior probabilities generated by CAVIARBF^75, 76^. The original GWAS summary statistics files were obtained from an atlas of genetic associations from the UK Biobank^77^. We manually fine-mapped the Chronic Kidney Disease, eGFR, Blood Urea Nitrogen, and Gout traits using the same code that was used to generate the CausalDB database (github.com/mulinlab/CAUSALdb-finemapping-pip). The summary statistics for all of these traits are available at the CKDGen Consortium site (ckdgen.imbi.uni-freiburg.de/)^78, 79^. We also manually fine-mapped the Hypertension trait and the original summary statistics can be found on the EBI GWAS Catalog^80^. We only looked at causal SNPs with a posterior causal probability of at least 0.05 in order to ensure SNPs with low causal probabilities did not cause false positive signals. Also, since g-chromVAR selects a semi- random set of peaks with similar average accessibility and GC content as background peaks, the method has an element of randomness. In order to ensure stable results, we ran g- chromVAR 20 times and averaged the results. Cluster/trait z-scores were plotted using ggheat (github.com/yanwu2014/swne).

To link causal SNPs to genes, we used functions outlined in the chromfunks repository (github.com/yanwu2014/chromfunks, /R/link_genes.R). This involved the identification of causal peaks for each cell type and trait (minimum peak Z score of 1, minimum peak posterior probability score of 0.025). Sites were then linked to genes based on co-accessibility (Cicero) with promoter regions, occurring within 3000 base pairs of a gene’s transcriptional start site (TSS). Only sites associated with genes detected as expressed in 10% of TAL nuclei/cells (snCv3/scCv3) were included. Motif enrichment within the causal SNP and TAL associated peaks was performed using the “FindMotifs” function in Seurat and only motifs for TFs expressed in 10% of TAL nuclei/cells (snCv3/scCv3) were included (**Supplementary Table 28**). For a TAL-associated ESRRB TF sub-network, peaks were linked to genes using Cicero, then subset to those associated with TAL (C-TAL, M-TAL) marker genes that were identified using the “Find[All]Markers” function in Seurat for subclass.l3 (p value < 0.05). TFs were then linked to gene-associated peaks based on the presence of the motif and correlation of peak and TFBS co-accessibility (chromVAR), using a correlation cutoff of 0.3. Only TFs with expression detected within 20% of TAL cells or nuclei (snCv3/scCv3) were included. Eigenvector centralities were then computed using igraph and the TF-to-gene network visualized using “PlotNetwork” in chromfunks.

### Patient cohorts used for clinical association analyses

Neptune^81^ (199 adult patients) and ERCB^44^ (111 patients) expression data were used as validation cohorts to determine the significance between patients with different levels of cell state gene expression. NEPTUNE (NCT01209000) is a multi-center (21 sites), prospective study of children and adults with proteinuria recruited at the time of first clinically indicated kidney biopsy (**Supplementary Table 30**). The study participants were followed prospectively, every 4 months for the first year, and then biannually thereafter for up to 5 years. At each study visit, medical history, medication use, and standard local laboratory test results were recorded, while blood and urine specimens were collected for central measurement of serum creatinine and urine protein/creatinine ratio (UPCR) and eGFR (mL/min/1.73m2). End stage kidney disease (ESKD) was defined as initiation of dialysis, receipt of kidney transplant or eGFR <15 mL/min/1.73m2 measured at two sequential clinical visits; and the composite endpoint of kidney functional loss by a combination of ESKD or 40% reduction in eGFR^82^. Genome wide transcriptome analysis was performed on the research core obtained at the time of a clinically- indicated biopsy using RNA-sequencing (RNA-seq) by the University of Michigan Advanced Genomics Core using Illumina HiSeq2000. Read counts were extracted from the fastq files using HTSeq (version 0.11). Neptune mRNA sequencing and clinical data are controlled access data and will be available to researchers upon request to.

ERCB is the european multicenter study that collects biopsy tissue for gene expression profiling across 28 sites. Transcriptional profiles of biopsies from patients in the ERCB were obtained from GEO (GSE104954).

### Clinical association of cell state scores

The gene expression data from tubulointerstitial compartment of the kidney biopsies from Neptune patients was used to calculate the composite scores for the genes enriched in degenerative, aPT, aTAL, and aStr states. The expression of the genes that were uniquely enriched in the cell state (described above) and that were found in both snCv3 and scCv3 were used to calculate the composite cell state score (**Supplementary Table 27**). Since scCv3 did not efficiently identify all stromal cell types, the union of the enriched genes from scCv3 and snCv3 data were used to calculate the aStr cell state score. We also generated a cell state score for the genes that were commonly enriched in aPT and aTAL cells.

For outcome analyses (40% loss of eGFR or ESKD), patient profiles were binned according to the degree of cell state score by tertile. Kaplan-Meier (K-M) analyses were performed using log rank tests to determine significance between patients with different levels of cell state gene expression. In the ERCB cohort, differential expression analyses were performed between the cell state scores in the disease group and living donors. The cell state scores for both Neptune and ERCB bulk mRNA transcriptomics data were generated^22^. Briefly, the cell state scores were generated by creating Z scores for each of the cell state gene sets and then using the average Z score as the cell state composite score.

### Sample level analysis and clustering on clinical association gene sets

To find association between the expression patterns of patients and clinical genesets (see previous method). We performed sample-level clustering using the expression profiles from the clinical genesets (**Supplementary Table 27**). All the cells from the same patient in snCv3 and scCv3 were aggregated to get pseudo-bulk count matrices. The matrices were further normalized by RPKM followed by tSNE dimension reduction. Groups of patients were then identified based on k-means clustering and density based methods in the reduced spaces. Patients identified as the same clusters were grouped together. To associate the patient pattern with clinical features, we calculated the distribution of eGFR in each identified group (see code repo).

To identify genesets that best differentiate AKI and CKD patients in Adaptive PT and TAL cell population, we trained a gene-specific logistic regression model based on the sample-level gene expression, the model was used to assess the predictive power that differentiate AKI and CKD patients in both snCv3 and scCv3 measured by area under the curve (AUC). The genes with AUC > 0.65 on both snCV3 and scCv3 were selected for downstream analysis (**Supplementary Table 29**).

### Pseudo-time analysis of PT and TAL cells

To find cells associated with disease progression, we performed trajectory analysis for PT and TAL cells. To get accurate pseudo-time and trajectory estimation, we removed degenerative cell populations in both PT and TAL and inferred the trajectory for single nuclei and single cell separately using the Slingshot package^83^ (Verson: 2.0.0). We specified normal cell populations as the start points for trajectory inference (S1-S3 in PT and M-TAL in TAL) using Slingshot parameter start.clus. The correspondent trajectory embedding was visualized using plotEmbedding function in the pagoda2 package.

To identify if the gene expression was statistically significantly associated with the inferred trajectory, we modeled the expression of a gene as a function of the estimated pseudo-time by fitting a gam model with cubic spline regression using formula expi ∼ t, where t is the pseudo-time. The model is then compared to a reduced model expi ∼ 1 to get p-value estimates using F-test. Benjamini-Hochberg method was used to calculate the adjusted p-values. To further identify the conditional differences of expression along the trajectory, we extended the base gam model by fitting a conditional-smooth interaction using “CKD” as a reference. The significant results for the extended model show the genesets whose expression levels are conditionally different along the inferred trajectory. We visualized the smoothed curve along with expression values for specific genes as a function of pseudo-time, which was implemented in plot_gene_psedotime function (see code repo).

### Gene module detection and cell assignment

To identify expression modules for significant gene sets along estimated trajectory, we applied the module detection algorithm implemented in WGCNA package^84^ (Version: 1.70-3) based on the smoothed gene expression matrix with parameters softPower=10 and minModuleSize=20. The similar modules detected by the original parameters were further merged. In total, we identified 5 different modules in PT and 6 modules in TAL cells. For the genesets in each module, we further performed pathway analysis using the Reactome online tool^85^ (reactome.org/PathwayBrowser/). In addition, to determine the direction of disease progression, we investigated the enrichment of clinical associated gene sets for each module by performing log ratio enrichment tests (**Extended Data Fig. 12c, g**).

To identify cells that are associated with each module, we developed a systematic approach. Briefly, for the cells in the smoothed expression matrix, we performed dimension reduction using PCA followed by louvain clustering. This allowed identification of cell clusters along the trajectory. For the identified cell clusters, we then did hierarchical clustering to calculate the correlation of each module based on mean gene expression values and further linked the clusters with associated modules by cutting the hierarchical tree. Finally, module labels for each cell were assigned based on its associated clusters. To link scCv3 data sets with snCv3 modules, we performed k-means clustering based on the joint embedding of PT/TAL cells and assigned the cells in scCv3 to modules based on the majority voting from its k’s nearest neighbors (see code repo).

To further investigate cluster-free compositional change between disease conditions, we also performed cell density analysis, where we compared the normalized cell density between AKI and CKD conditions through 2D kernel estimates using Cacoa Package (github.com/kharchenkolab/cacoa). Z-scores were calculated to identify the regions that showed significant differences of cell density.

### SLIDE-Seq2

#### Puck preparation and sequencing

Tissue pucks were prepared and sequenced^18, 86^ according to the step-by-step protocol: dx.doi.org/10.17504/protocols.io.bvv6n69e. Libraries were sequenced on a NovaSeq S2 flowcell (NovaSeq 6000) with a standard loading concentration of 2nM (read structure: Read 1 - 42 bp, Index 1 - 8 bp, Read 2 - 60 bp, Index 2 - 0 bp). Demultiplexing, genome alignment and spatial matching was performed using Slide-seq tools github.com/MacoskoLab/slideseq-tools/releases/tag/0.1.

#### Deconvolution

We used Giotto^87^ (version 1.0.3) for handling the slide-seq data and RCTD^88^ (version 1.1.0) for the cell type deconvolution. Since only reference tissue was used for slide- seq and it only contained the kidney cortex, all degenerative states and medullary subtypes were removed from the snCv3 cell subclasses prior to deconvolution. The counts from all beads across all pucks were pooled and deconvolved hierarchically: first, the broad subclass level 1 annotations in the Seurat object were used to deconvolve all beads (gene_cutoff = 0.0001, gene_cutoff_reg = 0.00015, fc_cutoff = 0.4, fc_cutoff_reg = 0.5). The prediction weights were normalized to sum to 100 per bead. Beads for which one cell type had a relative weight of 50% or higher were classified as that cell type. Then, for each level 1 subclass, all classified beads were further deconvolved using the level 2 annotation of that subclass, as well as the remaining subclass level 1 annotations (same parameters as level 1). Classification at subclass level 2 was done similar to level 1. Note that the bulk parameters in RCTD were fitted using all beads before subsetting the RCTD object to contain only beads confidently classified to a specific subclass. For all further analyses, we used only those pucks for which the median UMI per bead was higher than 100 (puck IDs with the format Puck_20090X_XX).

#### Cell type interaction

For each puck we first consolidated all the subclass level 2 immune subtypes, then subsetted to those beads that had a level 2 classification (relative weights greater than 50%). Delauney network was constructed for the remaining beads and Giotto’s “cellProximityEnrichment” was used to find the proximity enrichment of cell types at annotation level 2. To generate the interaction plot in **Figure 2d**, the enrichment values for each cell type pair were averaged across all pucks. The heterotypic interactions with enrichment higher than 0.6 were plotted with “cellProximityNetwork” in Giotto.

#### 10X Visium

Human kidney tissue was prepared and imaged according to Visium Spatial Gene Expression protocols (10x Genomics) according to the manufacturer protocol (CG000240 protocol, Visium Tissue Preparation Guide) and as previously described ^89^. Tissue was sectioned at 10 µm thickness from Optimal Cutting Temperature (OCT) compound embedded blocks. A Keyence BZ-X810 microscope equipped with a Nikon 10X CFI Plan Fluor objective was used to acquire hematoxylin and eosin (H&E) stained brightfield mosaics which were subsequently stitched. mRNA was isolated, libraries prepared, and sequencing was performed on an Illumina NovaSeq 6000^90^. mRNA was isolated from stained tissue sections after permeabilization for 12 minutes.

Released mRNA was bound to oligonucleotides in the fiducial capture areas. mRNA was then reverse transcribed and underwent second strand synthesis, denaturation, cDNA amplification, and SPRIselect cDNA cleanup (Visium CG000239 protocol). Space Ranger (v1.0.0) with the reference genome GRCh38 3.0.0 was used to perform expression analysis, mapping, counting, and clustering. Normalization was performed with SCTransform. Final data processing was done in Seurat (v3.2.3). A transfer score system was used to assess and map the proportion of signatures arising from each 55 µm spot. The transfer score reflects a probability between each spot’s signature and its association with a given snCv3 subclass (level 3). Seurat transfers the snCv3 subclass labels according to the transfer score. The highest probability transfer scores have the highest proportion mapped within each spatial transcriptomics spot pie graph. In cell state analyses, instead of mapping the subclasses, the six cell states annotated in snCv3 were mapped across all spots in the samples. To determine whether the 75 snCv3 subclasses (level 3) were appropriately mapped to histologic structures, the proportion of signature in each spot was compared to a histologically validated set of six unsupervised clusters defined by Space

Ranger (in **Extended Data Fig. 7D**)^89^. These six unsupervised clusters (glomerulus, proximal tubule, loop of Henle, distal convoluted tubule, connecting tubule and collecting duct, and the interstitium) had an overall alignment of 97.6% with the underlying histopathologic structures in the H&E image.

### Label-free and multi-fluorescence large-scale 3D imaging

Kidney biopsy cores frozen in OCT from patients with acute kidney injury or chronic kidney disease enrolled in KPMP were used for label-free imaging followed by multiplexed-fluorescence large scale 3D imaging as outlined in the following protocol: dx.doi.org/10.17504/protocols.io.9avh2e6, and described in a recent publication by Ferkowicz et al.^27^. Frozen biopsies were sectioned to a thickness of 50 µm using a cryostat and then immediately fixed in 4% fresh paraformaldehyde (PFA) for 24 hrs, and subsequently stored at 4°C in 0.25% PFA.

The first step in imaging consists of label-free imaging with multiphoton microscopy to collect autofluorescence and second harmonic images of the unlabeled tissue mounted in non-hardening mounting medium. Imaging was conducted using a Leica SP8 confocal scan-head mounted to an upright DM6000 microscope. For large-scale imaging of tissues at submicron resolution, the Leica Tile Scan function was used to collect a mosaic of smaller image volumes using a high-power, high-numerical aperture objective. Leica LASX software (v. 3.5) was then used to stitch these component volumes into a single image volume of the entire sample. The scanner zoom and focus motor control were set to provide voxel dimensions of 0.5 x 0.5 um laterally and 1 um axially.

Labeling of tissue for fluorescence microscopy was preceded by washing in phosphate-buffered saline (PBS) and blocking with PBS with 0.1% Triton X-100 (MP Biomedical) and 10% Normal Donkey Serum (Jackson Immuno Research). Antibodies for indirect immunofluorescence were applied first for 8-16 hours at room temperature, followed by washing cycles of PBS with 0.1% Triton X-100. Incubation cycle with secondary antibodies occurred next, followed by washing and finally application of directly labeled antibodies. Antibodies targeting markers for tubular cells and structures (Aquaporin-1, Uromodulin, F-actin) and immune cells (Myeloperoxidase, CD68, CD3, Siglec 8) were used, in addition to nuclei labeling using DAPI (**Supplementary Table 31**). After final washing cycles, the tissue was mounted in Prolong Glass (Thermo Fisher).

Confocal microscopy was conducted using a Leica 20x 0.75 NA multi-immersion objective (adjusted for oil immersion), with excitation sequentially provided by a solid state laser launch with laser lines at 405 nm, 488 nm, 552 nm and 635 nm. Images in 16 channels (emission spectra collected by PMT detectors adjusted for the following ranges: 410-430nm, 430-450nm, 450-470nm, 470-490 nm, 500-509nm, 510-519nm, 520-530nm, 530-540nm, 570-590nm, 590-610nm, 610-630nm, 631-651nm, 643-664nm, 664-685nm, 685-706nm and 706-726nm) were collected for each focal plane of each panel of the 3D mosaic. The resulting 16-channel image is then spectrally deconvolved (via linear unmixing using the Leica LASX linear unmixing software) to discriminate the 8 fluorescent probes in the sample. Validation of the linear unmixing has been described in a previous publication^27^.

### Confocal immunofluorescence microscopy

Human kidney tissue samples from cortex or medulla were fixed in 4% PFA, cryopreserved in 30% sucrose and frozen in O.C.T cryomolds, and were cut into 5 μm sections. Sections were post fixed with 4% PFA for 15 min at room temperature, blocked in blocking buffer (1% BSA, 0.2% skim milk, 0.3% Triton x-100 in 1X PBS) for 30 minutes at room temperature and then immunofluorescence microscopy was performed by first using overnight incubation at 4 ^0^C with primary antibodies and then followed by labeling with secondary antibodies. The primary antibodies included NRXN-1beta, Tuj1, collagen I & III, Synapsin-1, NPSH-1, SLC14A2, UMOD, CD31, CD34, CD11b, PROM1, KIM1, VCAM1, AQP1, AQP2, CD45 and S100 (**Supplementary Table 32**). After washing, labeling with the secondary antibodies was performed using Alexa- 488 conjugated goat anti-mouse IgG, or Cy3- conjugated goat anti-rabbit IgG, or Cy5- conjugated donkey anti-goat IgG at room temperature for one hour. After washing, sections were counterstained with DAPI for nuclear staining. Images were acquired with a Nikon 80i C1 confocal microscope.

### Tissue cytometry and in situ cell classification

Tissue cytometry and analysis were conducted using the Volumetric Tissue Exploration and Analysis (VTEA) software (v1.0a-r9). VTEA is a 3D image processing workspace that was developed as a plug-in for ImageJ/FIJI^91^. The version of VTEA which includes the supervised and unsuerives labeling of cells and combining spatial and features based gating strategies used here is available at: github.com/icbm-iupui/volumetric-tissue-exploration-analysis. In this analytical pipeline, each individual nucleus was segmented using an intensity thresholding and connected components segmentation built into VTEA and ImageJ. Each surveyed nucleus became a surrogate for its cell, to which the location and marker staining around or within the nucleus could be registered. This captured information could be used to classify cells based on marker intensity or spatial features using scatterplot displays that allow various gating strategies and statistical analysis, including export as .csv files of all segmented cells and the associated features^92^. Cells classified based on marker intensity are summarized in **Supplementary Table 33.** Gated cells were mapped back directly into the image volumes, which allowed immediate validation of the gates. In addition, direct gating on the image could be performed, which could trace all the cells within the chosen region-of-interest back to the data display on the scatter plot. Therefore, cell classification could also be performed based on direct annotation of regions-of-interest (ROIs) within the image volumes.

Using tissue cytometry, 14 cell classes were defined based on the following features:

• Proximal tubules (PT) cells: AQP1+ cells in cortex +/- brush border staining;
• Cortical thick ascending limbs cells, C-TAL: UMOD+ cells in cortex
• Glomerular cells (which encompass podocytes, glomerular endothelium and mesangial cells) annotated ROIs based on morphology and F-actin staining
• Cortical large and medium vessel cells: annotated ROIs based on morphology and F-actin staining.
• Cortical distal nephron cells (distal tubules (CD), connecting tubules (CNT) and collecting ducts (C-CD): AQP1-, UMOD- and annotated ROIs based on unique morphology in cortex.
• Medullary thick ascending limbs cells, M-TAL: UMOD+ cells in medulla
• Descending thin limbs cells (DTL): AQP1+ cells in medulla
• Medullary collecting ducts (M-CD): AQP1-, UMOD- and annotated ROIs based on unique morphology in medulla.
• Vascular bundles in the medulla (VB): annotated ROIs based on unique morphology in medulla and F-actin staining
• Neutrophils: MPO+ cells
• Activated macrophages: MPO-, CD68+ cells
• T cells: CD3+ cells
• Cells in altered regions: annotated ROIs based on loss of (unrecognizable) tubular morphology, expanded interstitium, increased fibrosis (by second harmonic generation imaging) and cell infiltrates.
• Not determined: unable to be classified based on the criteria above

Using such an approach,1,540,563 cells were classified from all the biopsies used in this analysis. Annotated ROIs were curated and vetted by the pixel wise agreement between 3 of 4 experts who performed the individual annotation on each biopsy specimen separately.

### 3D Neighborhood building and representation

3D neighborhoods were calculated for every cell in each biopsy using VTEA and a radius of 25 um (50 voxels in x and y and 25 voxels in z). For each 3D neighborhood, VTEA was used to calculate the features: fraction-of-total and sum of each labeled cell was calculated by VTEA. A list of neighborhoods, positions in 3D and their features was exported by biopsy specimen image as .csv files.

### Neighborhood visualization and statistical analysis

CSV files generated in VTEA for neighborhoods by biopsy specimen were imported into R (v 4.0.4), parsed for the features sum of each label and monotypic neighborhood removed. These features were scaled by Z-standardization and used for louvain community detection (R packages: FNN and igraph) and *t*-SNE manifold projection (R package: Rtsne). To understand the interactions within neighborhoods, pairwise interactions by neighborhood were tallied and plotted on a chord plot (R package: circlize) and Pearson’s correlation coefficients were calculated and plotted (R package: Hmisc and corrplot). Subclasses of neighborhoods, those with at least one cell with a specific label were selected and plotted as network plots (R package: igraph) with edges in CD3 and Altered neighborhoods scaled at 40% of all other subclasses to facilitate visualization. All scripts are provided as an annotated RStudio notebook file(.Rmd).

### Data Availability

Raw sequencing and imaging data (snCv3, scCv3, 3D imaging) generated as part of the Kidney Precision Medicine Project (KPMP) has been deposited at atlas.kpmp.org. Raw sequencing data (snCv3, SNARE2, Slide-seq) generated as part of the Human Biomolecular Atlas Project (HuBMAP) has been deposited at portal.hubmapconsortium.org/. Raw sequencing data (scCv3) on living donor biopsies as part of the Chan Zuckerberg Initiative (CZI) and Human Cell Atlas (HCA) will be available in the Gene Expression Omnibus (GEO) as GSE169285. Visium spatial transcriptomic data is available in GEO as GSE171406. Neptune sequencing and clinical data is available upon request to. ERCB data was obtained from GEO as GSE104954. KPMP snCv3 and scCv3 cell types and expression profiles can be interrogated using the KPMP Data Atlas Explorer: https://atlas.kpmp.org/explorer. snCv3 healthy reference data is available for reference-based single cell mapping by the Azimuth tool: azimuth.hubmapconsortium.org/.

### Code Availability

Code to reproduce figures will be available to download from github.com/KPMP/Cell-State- Atlas-2021.

## Acknowledgements

We are deeply indebted to the generosity of patients volunteering to donate tissue primarily for research purposes despite no direct immediate benefit to their clinical care. We thank the KPMP clinical coordinators at the recruitment sites for their efforts in patient enrolments and biopsy tissue procurement, pathologists, and the Central Hub for coordinating data collection, storage and making it accessible to the public through the consortium website. We thank the KPMP nomenclature working group for establishing the nephron schema and standardized nomenclature used, specifically Todd Valerius, Wilhelm Kriz, Brigitte Kaissling and Michael Rose. We are also grateful to the HuBMAP HIVE for building the infrastructure for data storage and public access to reference samples, in particular Jonathan Silverstein, Peter Kant, Katy Borner and Nils Gehlenborg. We thank the Indiana Center for Biological Microscopy for imaging assistance, Indiana Center for medical Genomics for sequencing of the 10X Visium samples and the Indiana Biobank for hosting the BBCI. We are grateful to the Kidney Translational Research Center (KTRC) at the Washington University (Division of Nephrology) and Mid America Transplant in St. Louis for infrastructural support for HuBMAP samples and Steve Steffan family in support of new omics technologies in kidney research (B1401-40 to SJ). This publication is part of the Human Cell Atlas - humancellatlas.org/publications.

The snCv3 and SNARE2 sequencing data were generated at the UC San Diego IGM Genomics Center supported by the National Institutes of Health (SIG grant #S10 OD026929) and Washington University Genome Technology Access Center at the McDonnell Genome Institute partially supported by NCI Cancer Center Support (#P30 CA91842) to the Siteman Cancer Center and by ICTS/CTSA (# UL1TR002345) from the National Center for Research Resources (NCRR). The KPMP data presented here is funded by the following grants from the NIDDK: U2C DK114886, UH3DK114861, UH3DK114866, UH3DK114870, UH3DK114908, UH3DK114915, UH3DK114926, UH3DK114907, UH3DK114923 and UH3DK114933. The HuBMAP data presented here is supported by U54HL145608 (SJ and KZ). Additional NIH support was provided by NIH/NIDDK K08DK107864 (M.T.E.); Indiana Grand Challenge Precision Health fund (R.M.F.); R01DK111651 (TME); P30 DK079312 (TME and PCD); U2CDK114886, U54DK083912, P30 DK081943 / HCA: Kidney Seed Network (MKZ); K23DK125529 (AN); U54DK083912 (SE); U01MH114828 (KKa); UH3CA246632 (EZM). The content is solely the responsibility of the authors and does not necessarily represent the official views of the National Institutes of Health.

## Author contributions

Coordination of manuscript writing and project: B.B.L., S.J. Patient Recruitment and Tissue Collection: A.K., A.S.N., C.P., D.S., E.H.K., F.P.W., J.C.W., J.R.S., M.K., P.M.P., R.D.T., S.J., S.R., S.S.W. Tissue Processing: A.K., A.S.N., D.B., D.S., E.A.O., J.R.S., M.K., M.T.E., P.C.D., S.J., S.R., S.W., T.M.E. RNA data generation: A.S.N., B.B.L., D.D., E.A.O., E.M., E.Z.M., F.C., K.S.B., K.Z., M.K., M.T.E., N.P., P.C.D., R.M., S.J., S.U., T.M.E. Imaging data generation: B.Z., D.B., M.T.E., P.C.D., S.J., S.W., T.M.E. ATAC data generation: B.B.L., D.D., K.Z., N.P., S.J. Data archive / infrastructure: B.B.L., D.D., M.K., M.T.E., P.C.D., Q.H., R.M., R.M.F., S.W., T.M.E., X.W. Data analysis: B.B.L., D.B., E.A.O., J.P.G., K.Ka., K.Z., M.K., M.T.E., P.C.D., P.V.K., Q.H., R.M., R.M.F., S.E., S.J., S.W., T.M.E., X.W., Y.W. Data interpretation: A.S.N., A.V., B.B.L., E.A.O., J.P.G., K.Ka., K.Z., M.K., M.T.E., P.C.D., P.H., .V.K., Q.H., R.M., R.M.F., S.E., S.J., S.W., T.M.E. Writing manuscript: B.B.L., K.Ka., K.Z., M.K., M.T.E., P.C.D., P.V.K., Q.H., R.M., R.M.F., S.J., S.W., T.M.E.

## List of collaborators

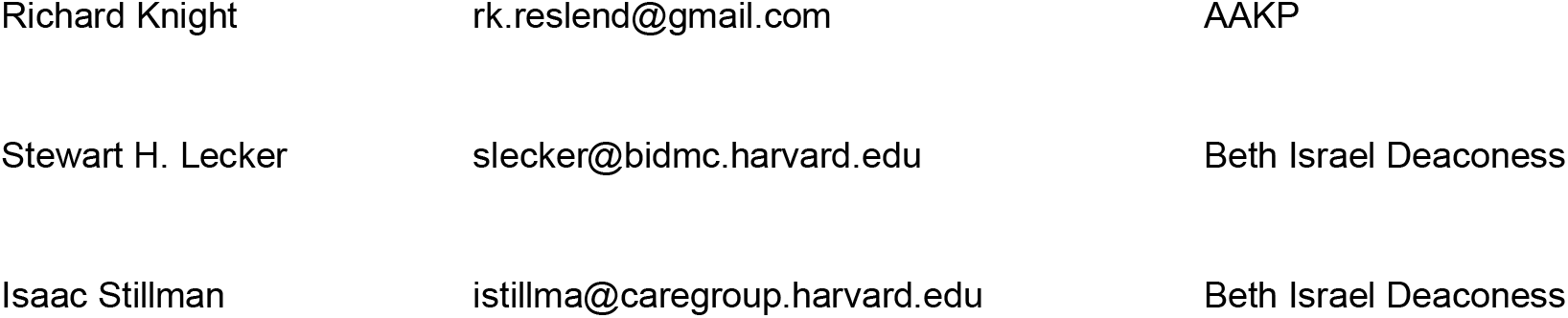
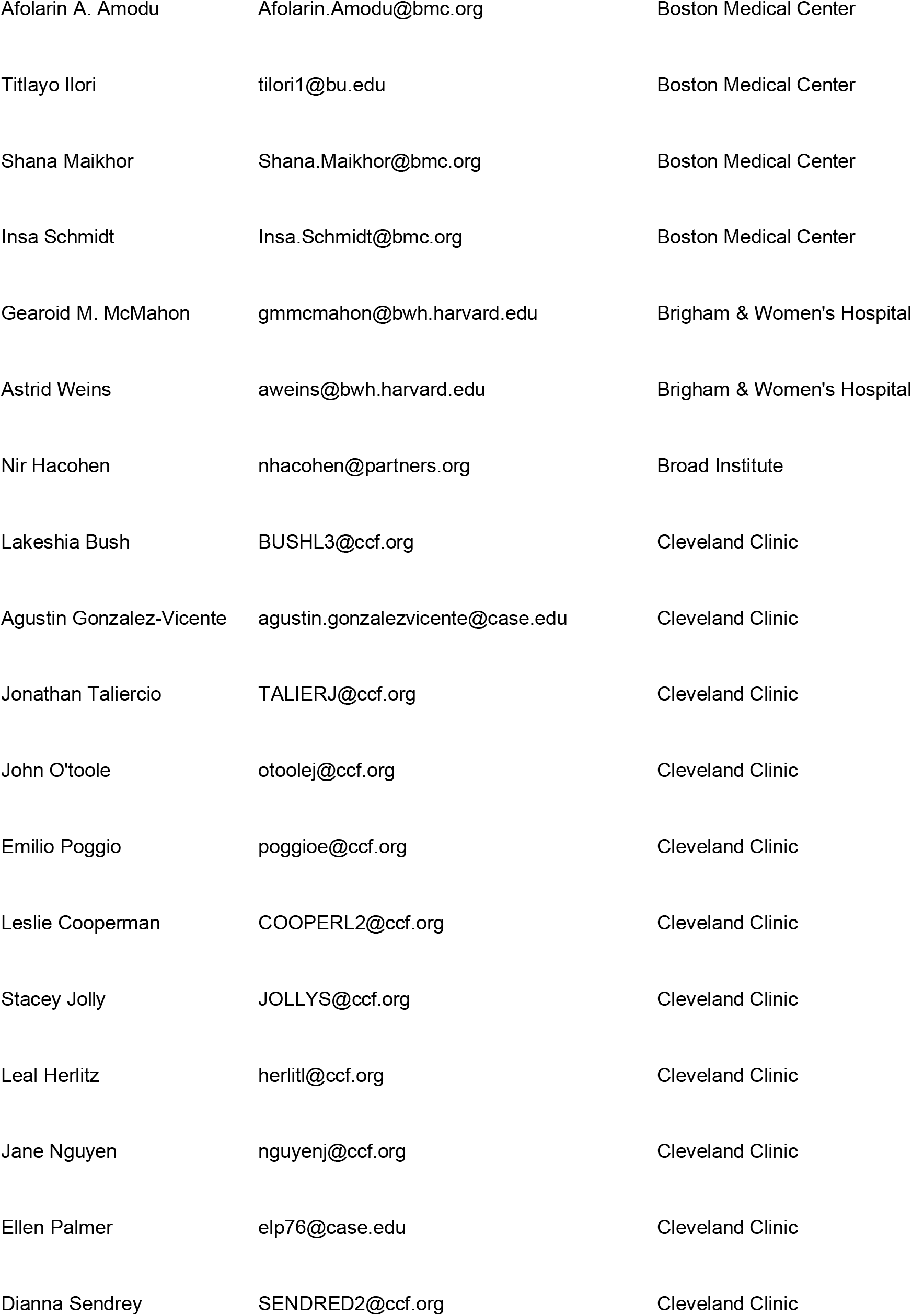
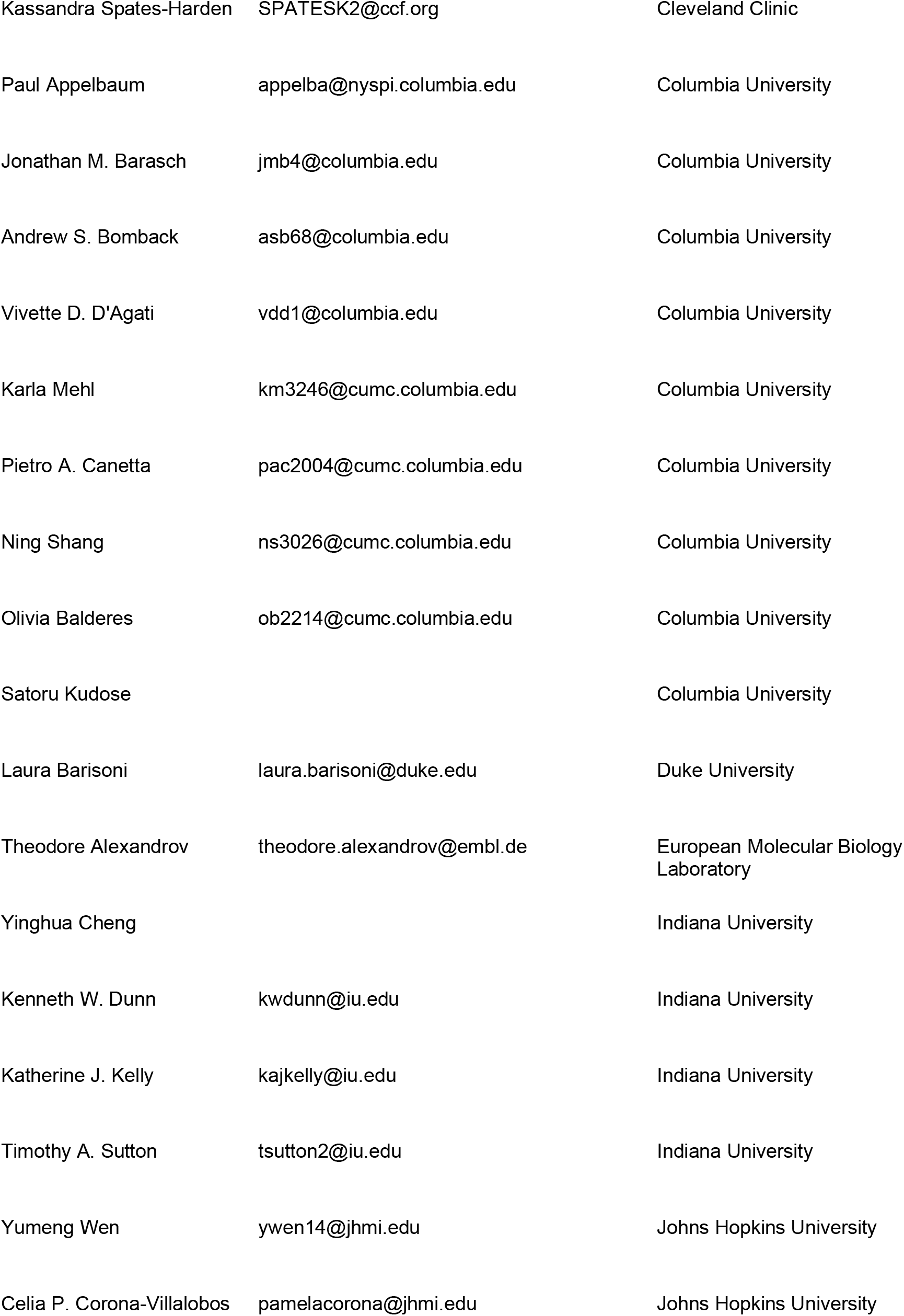
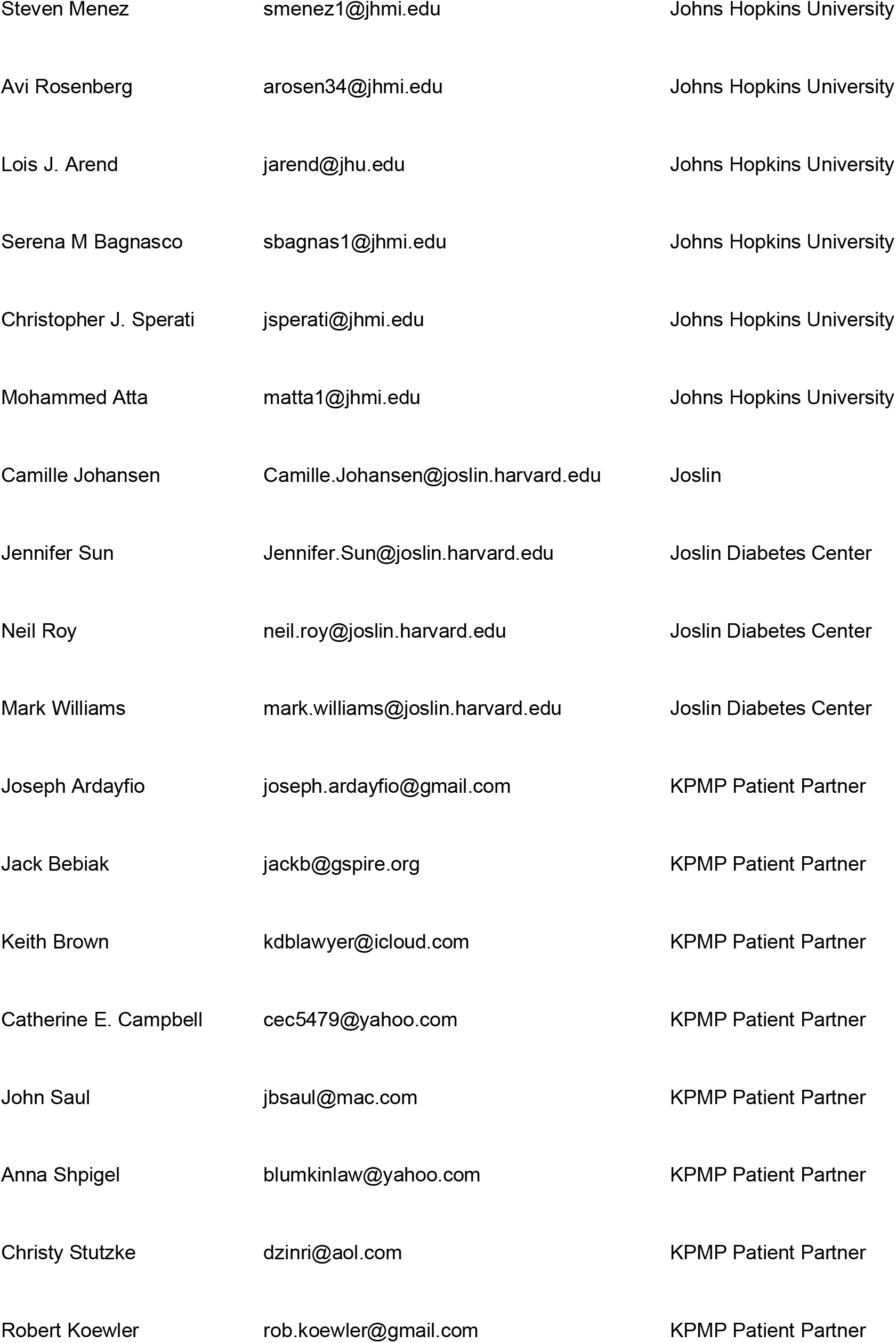
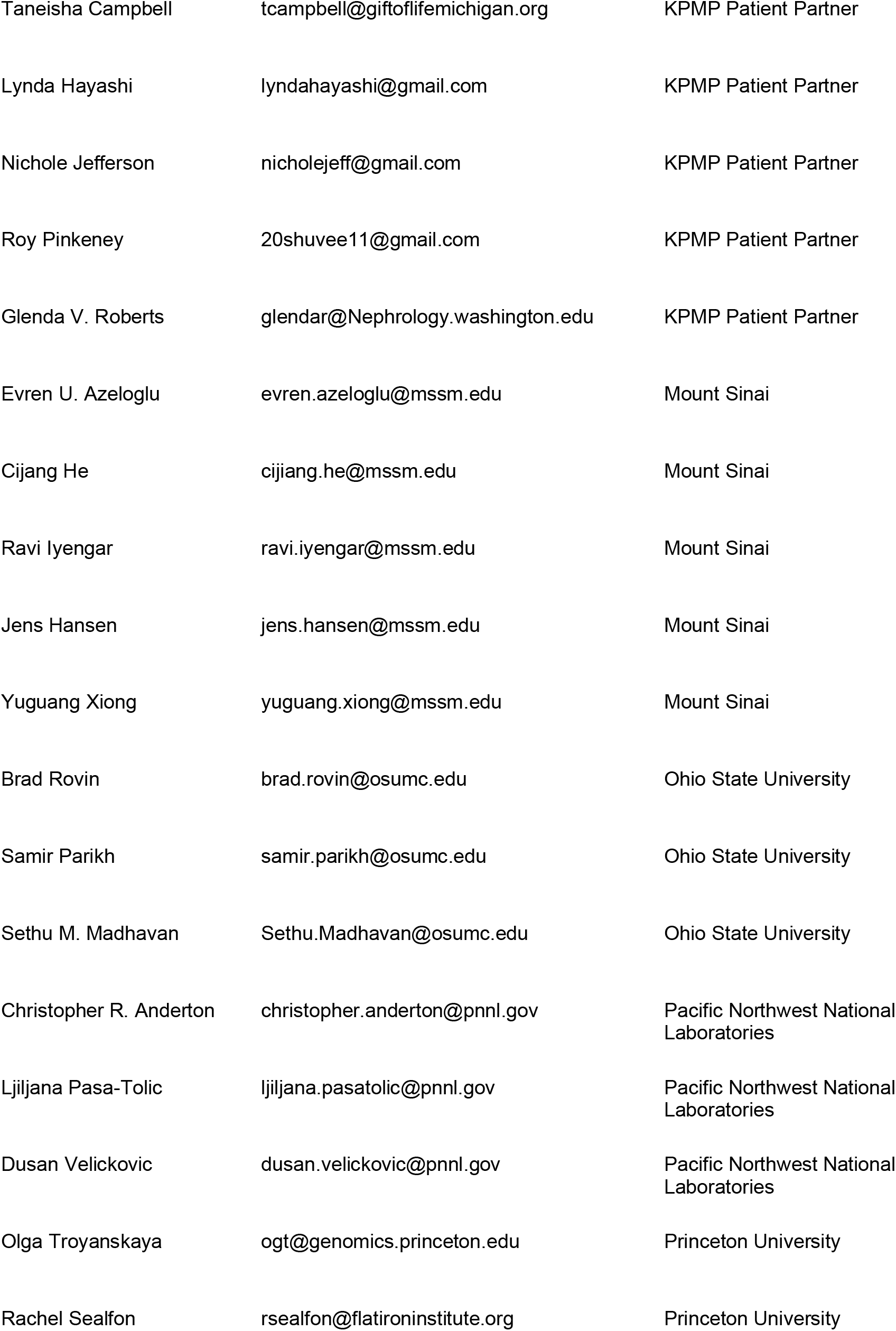
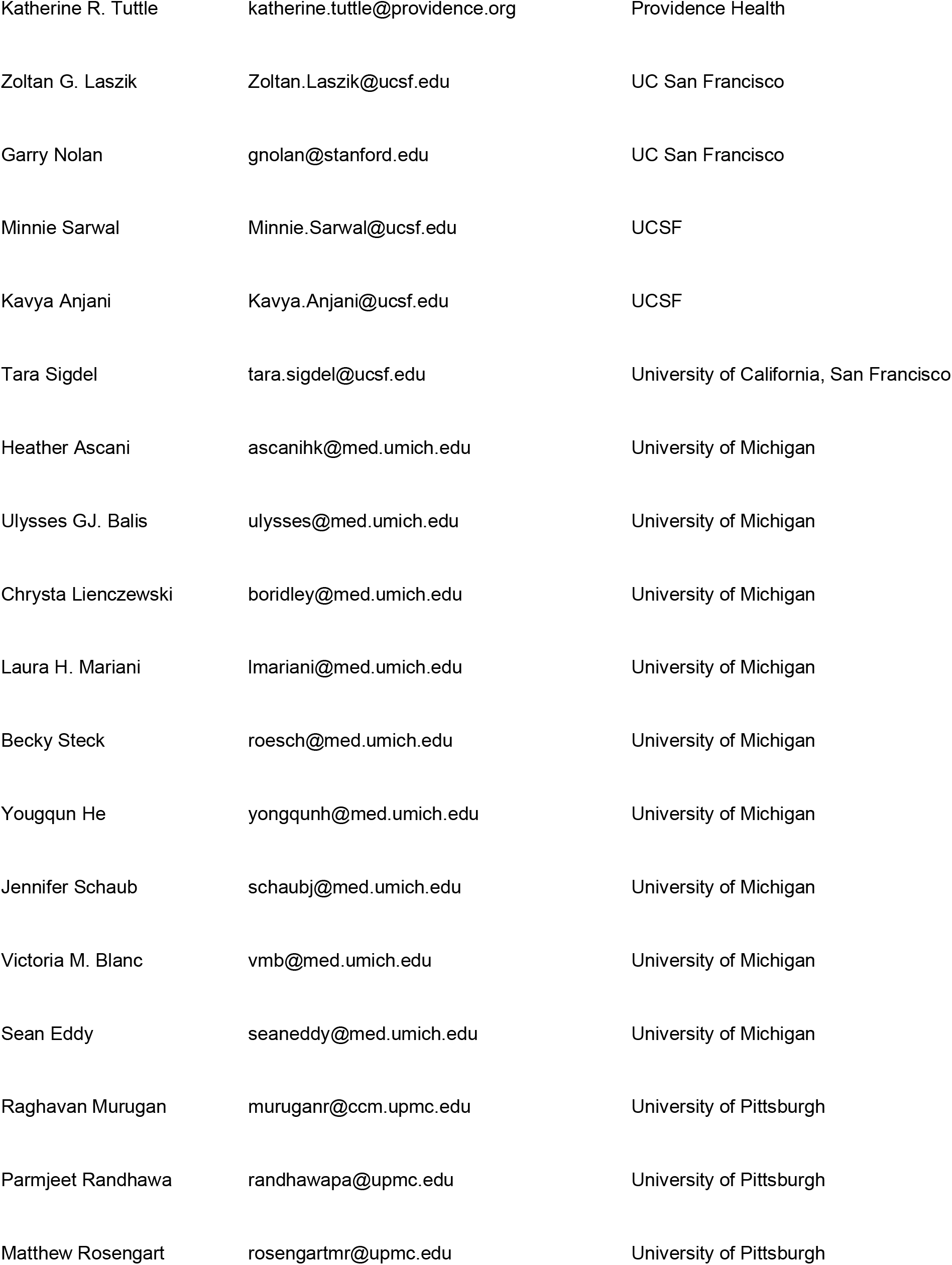
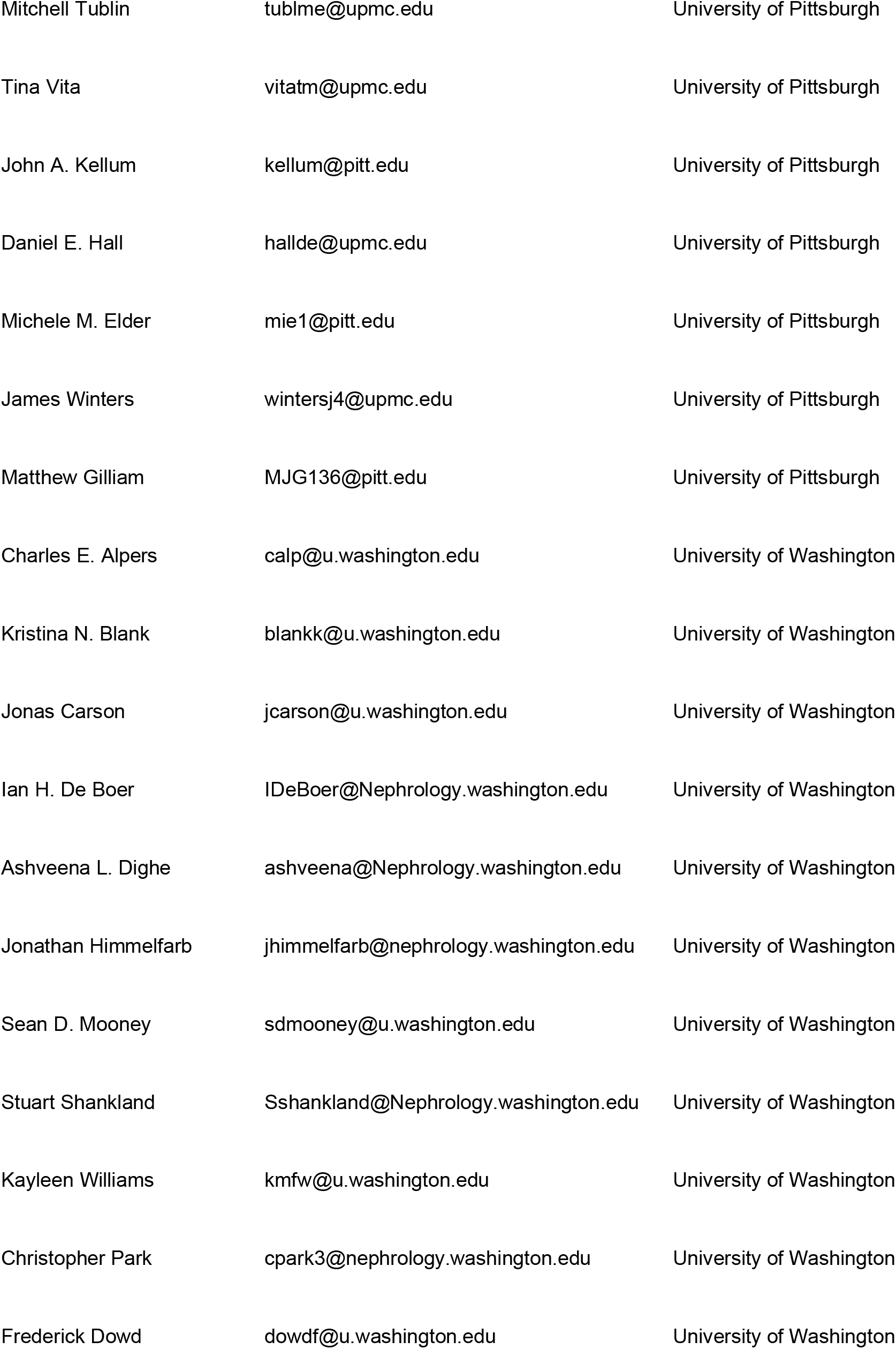
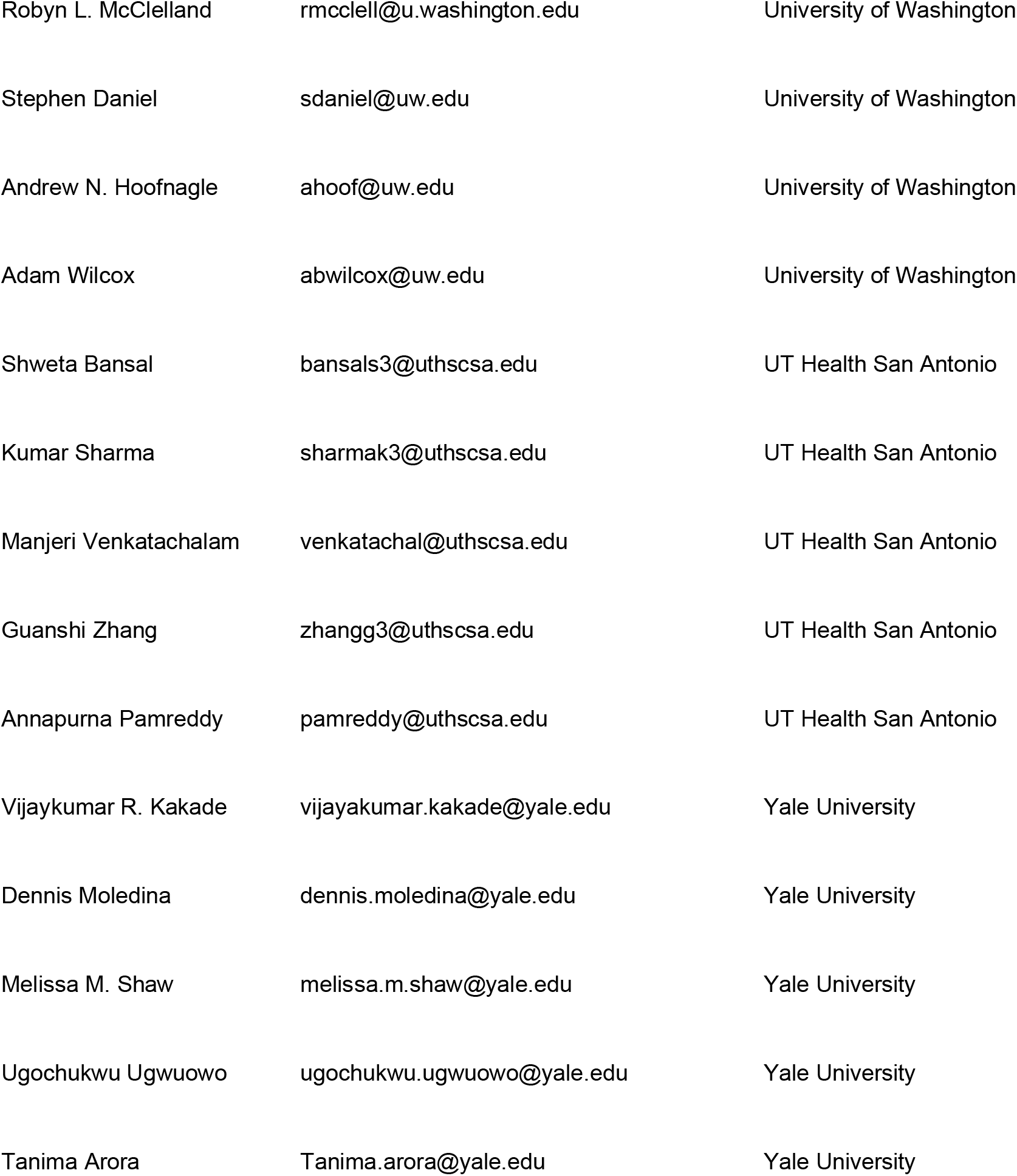

## Competing interests

No competing interests for the work submitted. Disclosures: P.V.K. serves on the Scientific Advisory Board to Celsius Therapeutics Inc. and Biomage Inc. A.V. is a consultant for Astute and NxStage; C.P. is a member of the advisory board of and owns equity in RenalytixAI, and serves as a consultant for Genfit and Novartis; M.K. has grants from JDRF, Astra-Zeneca, NovoNordisc, Eli Lilly, Gilead, Goldfinch Bio, Janssen, Boehringer-Ingelheim, Moderna, European Union Innovative Medicine Initiative, Chan Zuckerberg Initiative, Certa, Chinook, amfAR, Angion Pharmaceuticals, RenalytixAI, Travere Therapeutics, Regeneron, IONIS Pharmaceuticals, Astellas, Poxel, and a patent PCT/EP2014/073413 “Biomarkers and methods for progression prediction for chronic kidney disease” licensed; F.C. and E.Z.M. are paid consultants for Atlas Bio; P.M.P. is a consultant for Janssen; S.R. has research funding from AstraZeneca and Bayer Healthcare; J.R.S. consults for Maze, Goldfinch, and receives royalties from Sanfi Genzyme; K.Z. is a co-founder, equity holder and serves on the Scientific Advisory Board of Singlera Genomics. A.S.N. is on the external advisory board for CareDX. S.J. is a paid Blue SKy mentor for Meharry Medical College, Nashville and receives royalties from Elsevier Inc.

---

**Extended Data Figure 1.**
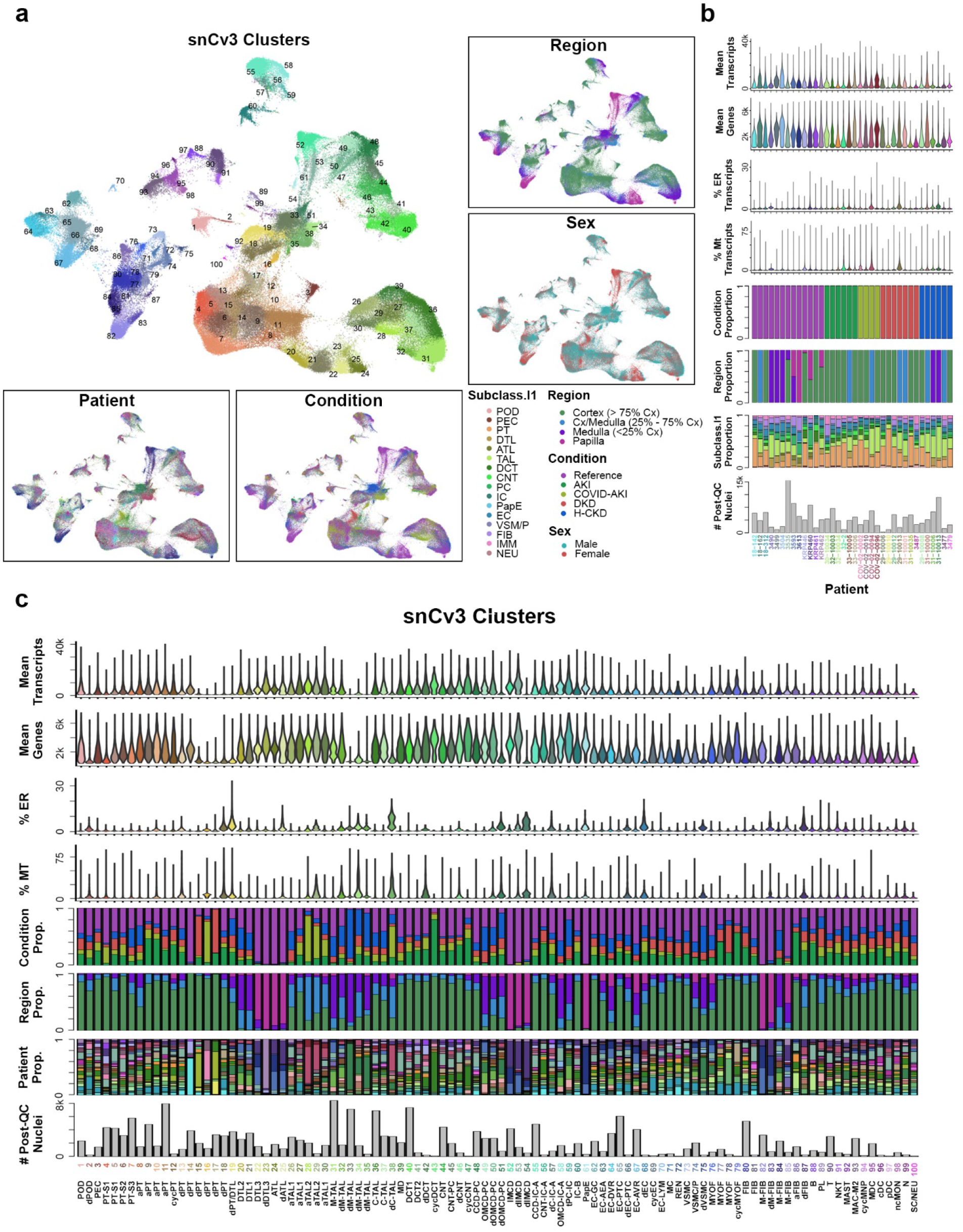
snCv3 cell types and quality metrics. **a.** UMAP plots for snCv3 clusters, with insets showing the corresponding tissue regions, sex, patient identities and conditions. **b.** Bar and violin plots for snCv3 patients shown in (**a**). Barplots showing the total number of post-QC nuclei used in the snCv3 clustering analysis, and the proportions that were associated with level 1 subclasses, regions sampled or the health or disease conditions. Violin plots show the percentage of transcripts associated with the mitochondria (Mt) or endoplasmic reticulum (ER), as well as mean genes and mean transcripts detected per patient sample. **c.** Bar and violin plots as in (**b**) for snCv3 clusters shown in (**a**), including proportion of nuclei contributed by each patient.

**Extended Data Figure 2.**
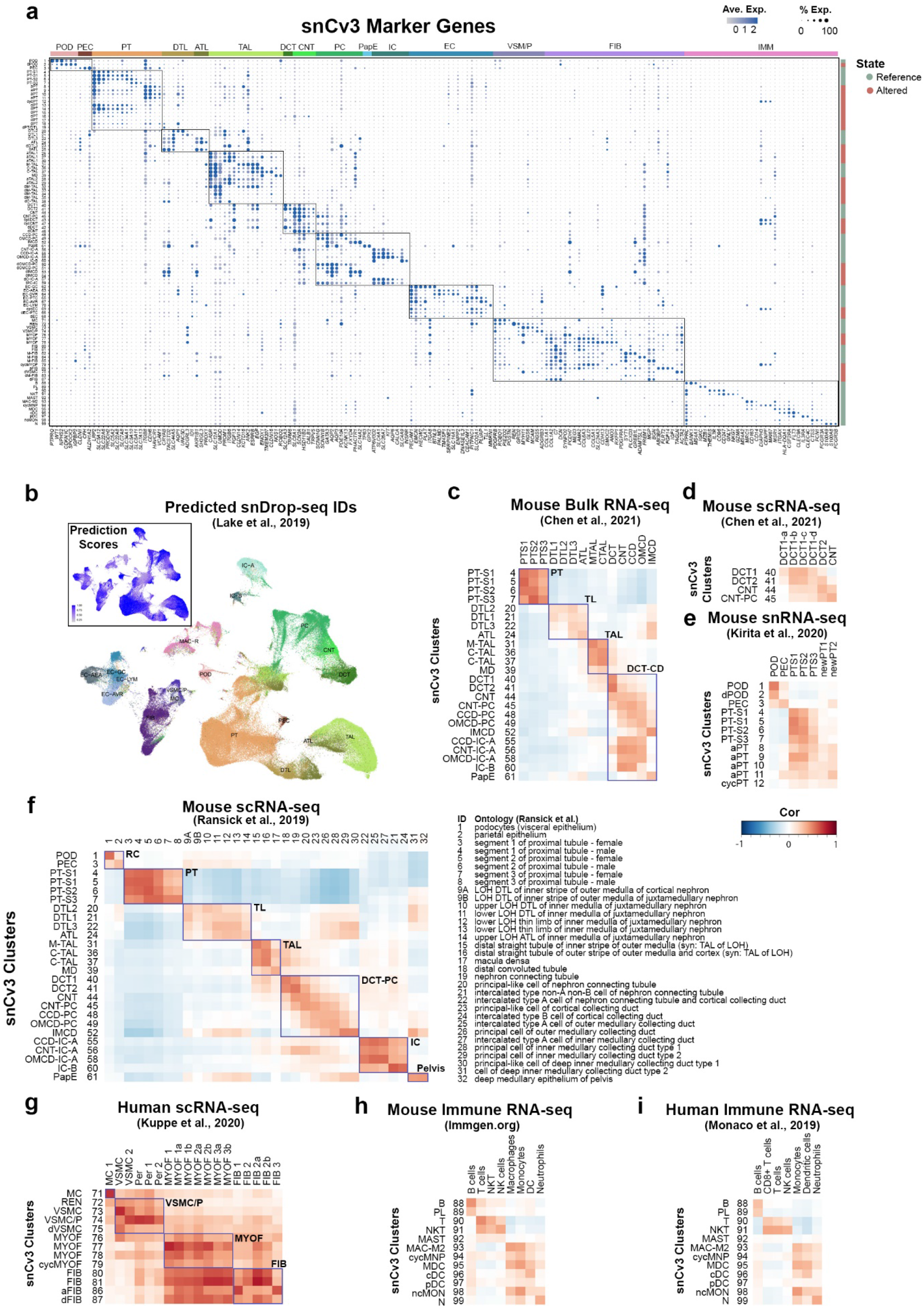
snCv3 marker genes and comparison with reference data. **a.** Dot plot showing averaged marker gene expression values (log scale) and proportion expressed for snCv3 clusters. **b.** Cell type labels predicted from Lake et. al. 2019^11^ mapped on the snCv3 UMAP embedding. Inset shows the corresponding prediction score values. **c.** Heatmap showing correlation of averaged scaled gene expression values for snCv3 epithelial (reference state) clusters and mouse bulk segmental RNA-seq data from Chen et al., 2021^58^. **d.** Heatmap showing correlation of averaged scaled gene expression values for snCv3 distal tubule clusters (reference states) and mouse scRNA-seq data from Chen et al., 2021^58^. **e.** Heatmap showing correlation of averaged scaled gene expression values for snCv3 clusters (reference and altered/adaptive states) and mouse snRNA-seq clusters from Kirita et al., 2020^3^. **f.** Heatmap showing correlation of averaged scaled gene expression values (reference states) for snCv3 clusters and mouse scRNA-seq clusters from Ransick et al., 2019^56^. **g.** Heatmap showing correlation of averaged scaled gene expression values for snCv3 stromal clusters (reference and altered/adaptive states) against human scRNA-seq clusters from Kuppe et al., 2020^23^. **h.** Heatmap showing correlation of averaged scaled gene expression values for snCv3 immune cell clusters and mouse immune cell types from Immgen.org. **i.** Heatmap showing correlation of averaged scaled gene expression values for snCv3 immune cell clusters and human immune cell types from Monaco et al. 2019^59^.

**Extended Data Figure 3.**
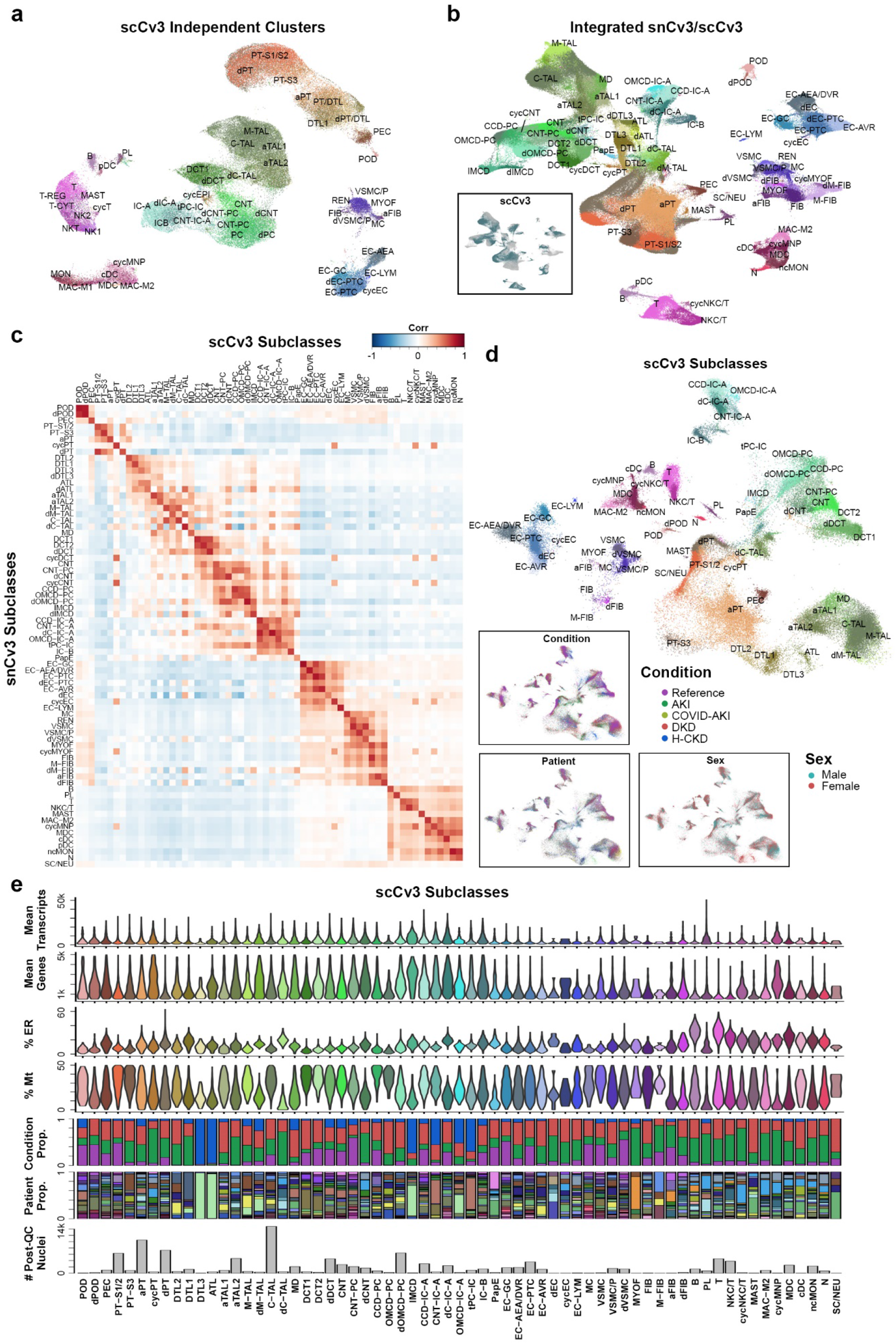
scCv3 integration and quality metrics. **a.** UMAP plot showing independent clustering and annotation of scCv3 data. **b.** UMAP showing integrated snCv3 and scCv3 clustering and harmonized subclass level 3 annotations. Inset shows location of scCv3 cells. **c.** Heatmap showing correlation of averaged scaled gene expression values for snCv3 and scCv3 using harmonized subclass level 3 annotations. **d.** UMAP plot showing scCv3 data projected into the snCv3 embedding shown in **Fig. 2b**. Insets show mapping of the corresponding sex, patient identities and conditions. **e.** Barplots showing the total number of post-QC nuclei per scCv3 subclass level 3, and the proportions that were associated with patients sampled or health/disease conditions. Violin plots show the percentage of transcripts associated with the mitochondria (Mt) or endoplasmic reticulum (ER), as well as mean genes and mean transcripts detected per subclass.

**Extended Data Figure 4.**
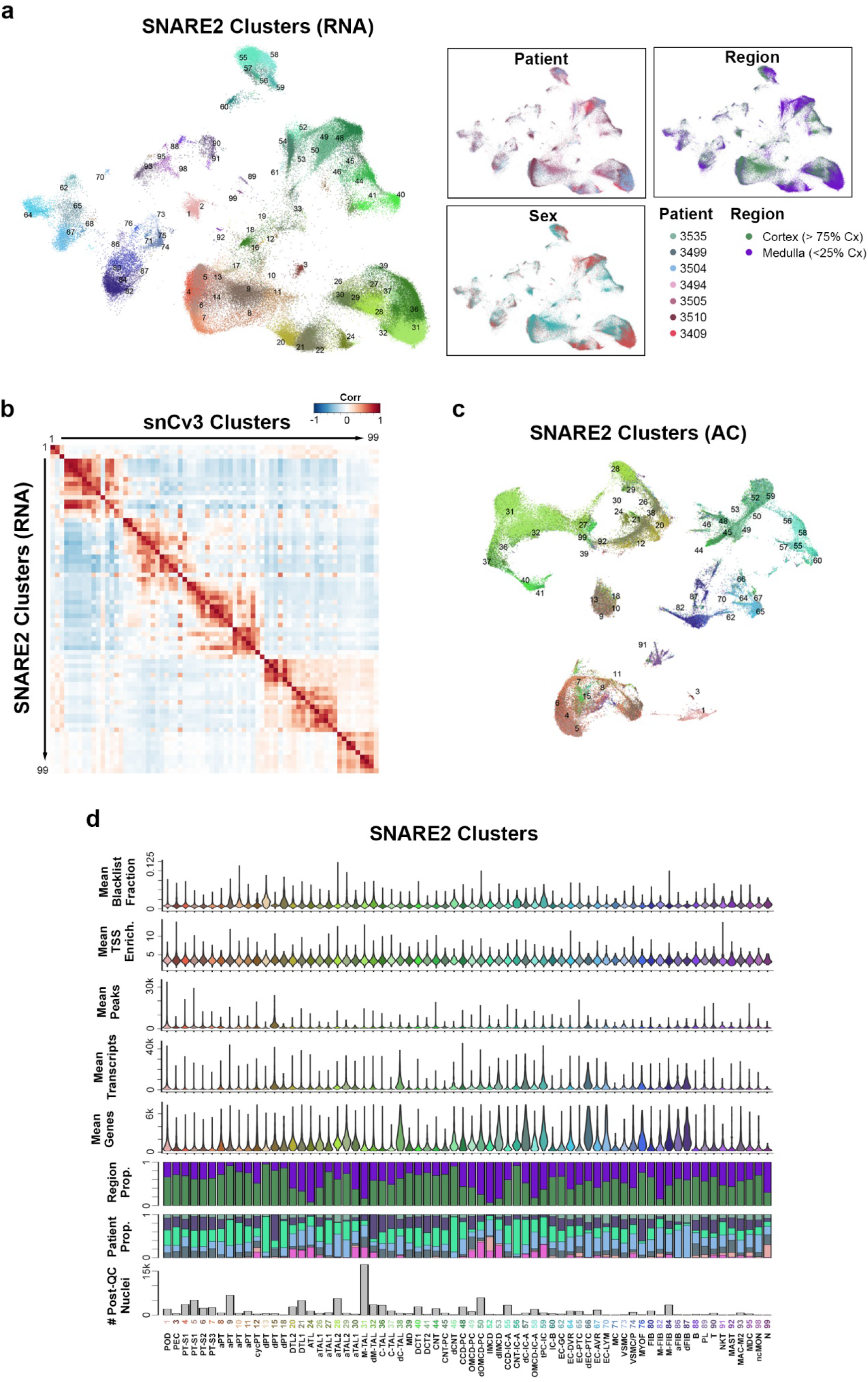
SNARE2 integration and quality metrics. **a.** UMAP plot showing SNARE2 RNA data projected onto the snCv3 embedding (**Fig. 2b**) and the corresponding harmonized cluster annotations. Insets show mapping of the tissue region, sex and patient identities. **b.** Heatmap showing correlation of averaged scaled gene expression values for SNARE2 and snCv3 using harmonized cluster annotations. **c.** UMAP embedding for SNARE2 AC based on Cistopic^70^ derived embeddings and showing harmonized clusters annotations as in (**a**). **d.** Barplots showing the total number of post-QC nuclei per SNARE2 cluster, and the proportions that were associated with patient or region sampled. Violin plots show the mean genes, transcripts (SNARE2 RNA) and mean peaks, TSS enrichments and blacklist fractions (SNARE2 AC) detected per cluster.

**Extended Data Figure 5.**
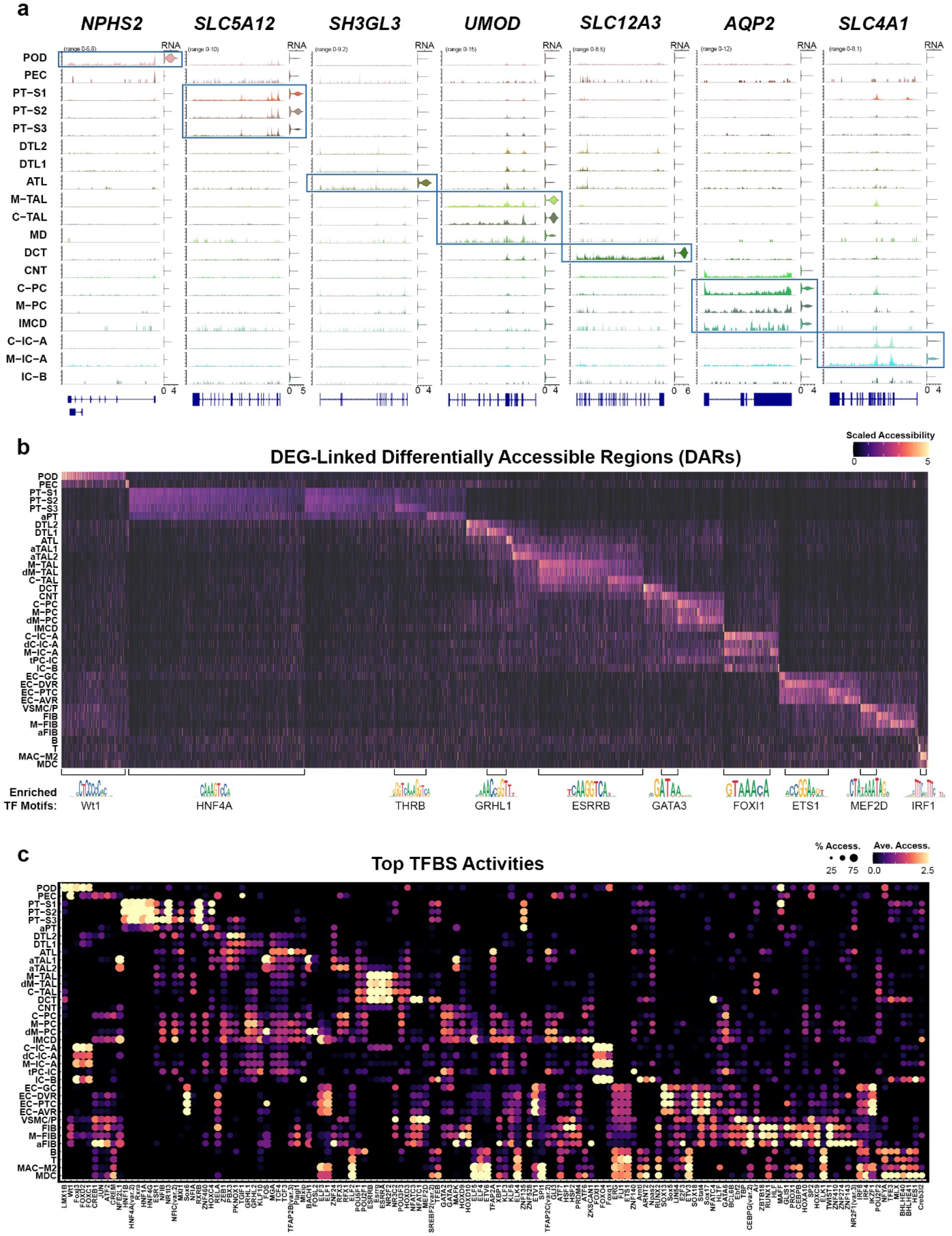
SNARE2 cell type regulatory profiles. **a.** Coverage plots showing SNARE2 AC read pile-ups for genomic regions associated with cell type marker genes. Violin plots show corresponding SNARE2 RNA gene expression values. **b.** Heatmaps showing averaged scaled chromatin accessibility values for differentially accessible regions (DARs) identified for cell type specific differentially expressed genes (DEGs, Methods). Select TF motifs enriched within the cell type specific DARs are shown. **c.** Dot plots showing average TFBS accessibilities (chromVAR) and proportion accessible for SNARE2 AC cell types.

**Extended Data Figure 6.**
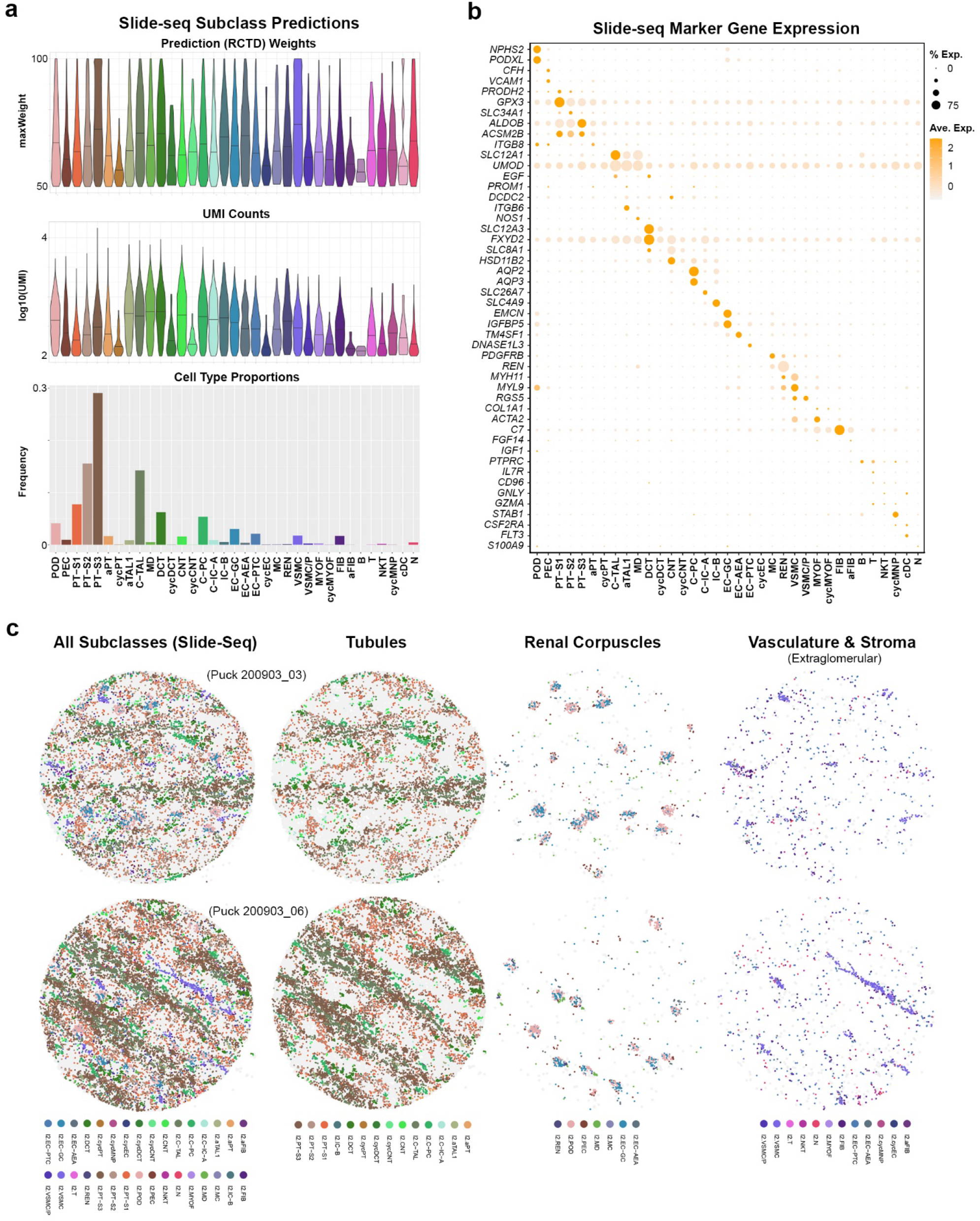
Slide-seq predicted cell types. **a.** Top: normalized RCTD weights for the beads classified at subclass level 2 (Methods). Middle: UMI counts per bead for classified beads. Bottom: relative frequency of cell types predicted across pucks. **b.** Expression of cell type markers identified by snCv3 in the classified Slide-seq beads. **c.** Two representative pucks showing subclass level 2 classifications. Cell types are grouped into 3 categories and plotted separately for clarity. For panels **a** and **b,** all pucks from a single individual with median UMI of 100 or more were pooled together. Puck diameter is 3mm.

**Extended Data Figure 7.**
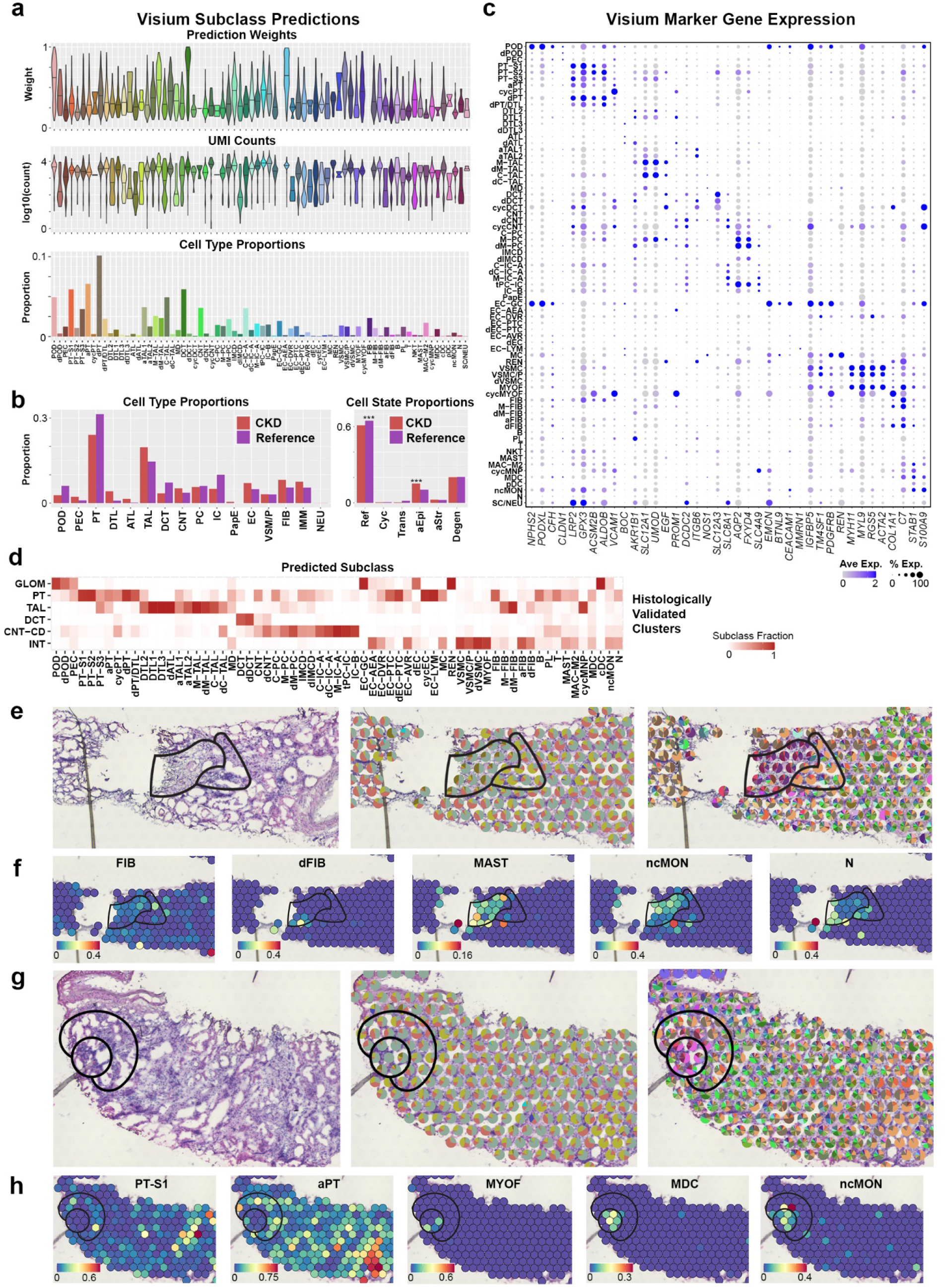
10X Visium predicted cell types. **a.** Analysis of subclass prediction on Visium spots for 4 reference nephrectomies and 4 biopsy specimens with chronic kidney disease (CKD). The top panel presents the distribution of transfer scores for the subclass (level 3) with the highest score in each spot. The middle panel presents the UMI counts associated with these spots. The bottom panel depicts the proportion of transcriptomic signatures for each subclass. In every spot subclass which had a non-zero transfer score, a fraction of the spot was assigned to the subclass, proportional to its transfer score relative to all non-zero transfer scores in that spot. **b.** Proportion of transcriptomic signatures in 4 CKD biopsies and 4 Reference nephrectomies. Left panel presents cell type classes and the right panel presents cell states. Where significance is indicated, p values are lower than 10^-4^ as calculated by a Fisher’s Exact test. **c.** Gene expression of select cell markers by predicted subclass (level 3) for all 8 samples. **d.** Alignment between the predicted cell type subclass and unsupervised clusters that were histologically validated (Methods). **e.** Detailed region of a CKD biopsy with fibrosis (left outline) and surrounding altered PT (right outline). The first panel presents the histological image, the middle panel shows the proportion of each cell state mapped to the spots, and the right panel shows the proportion of cell type subclasses. Each spot is 55 μm in diameter. **f.** Predicted transfer scores of fibroblasts and immune cell types in the region shown in (**e**). **g.** Detail of a region of immune cell infiltration (circle outline) and surrounding altered PT (outer crescent outline) on a CKD. From left to right: the histological image, the proportion of cell states predicted to each spot, and the proportion of subclasses. **h.** Predicted transfer scores for proximal tubules and MyoF and monocytes in the regions shown in (**g**).

**Extended Data Figure 8.**
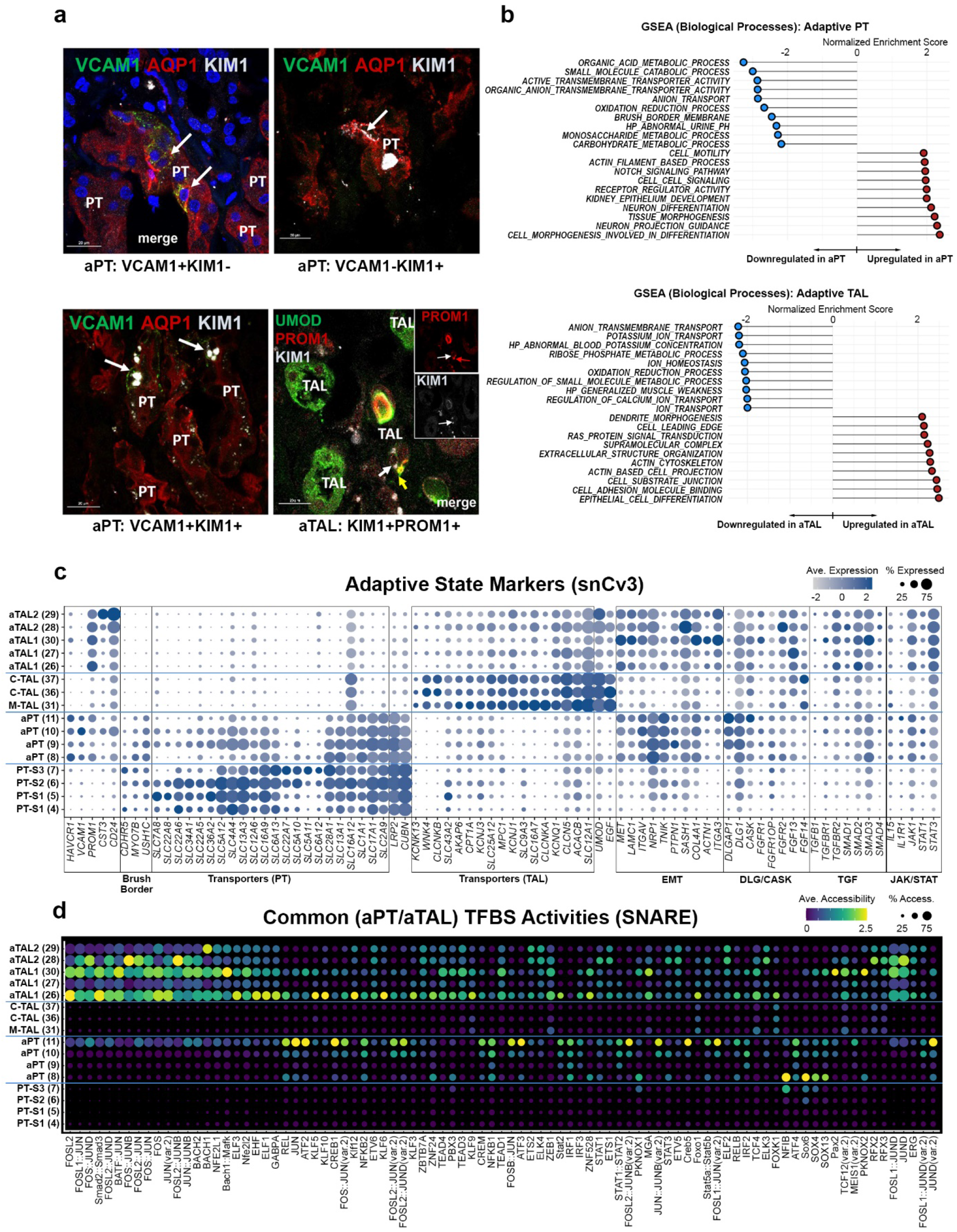
Adaptive epithelial state signatures. **a.** Immunofluorescent staining of VCAM1, AQP1, KIM1 (HAVCR1) in the aPT and UMOD, PROM1 and KIM1 in the TAL. Scale bars represent 20 µm. **b.** Gene Set Enrichment Analysis (GSEA) for genes upregulated or downregulated in adaptive epithelial states compared to reference states. **c.** Dot plot showing averaged marker gene expression values (log scale) and proportion expressed for snCv3 clusters. **d.** Dot plots showing SNARE2 average accessibilities (chromVAR) and proportion accessible for TFBSs showing differential activity in both aPT and aTAL.

**Extended Data Figure 9.**
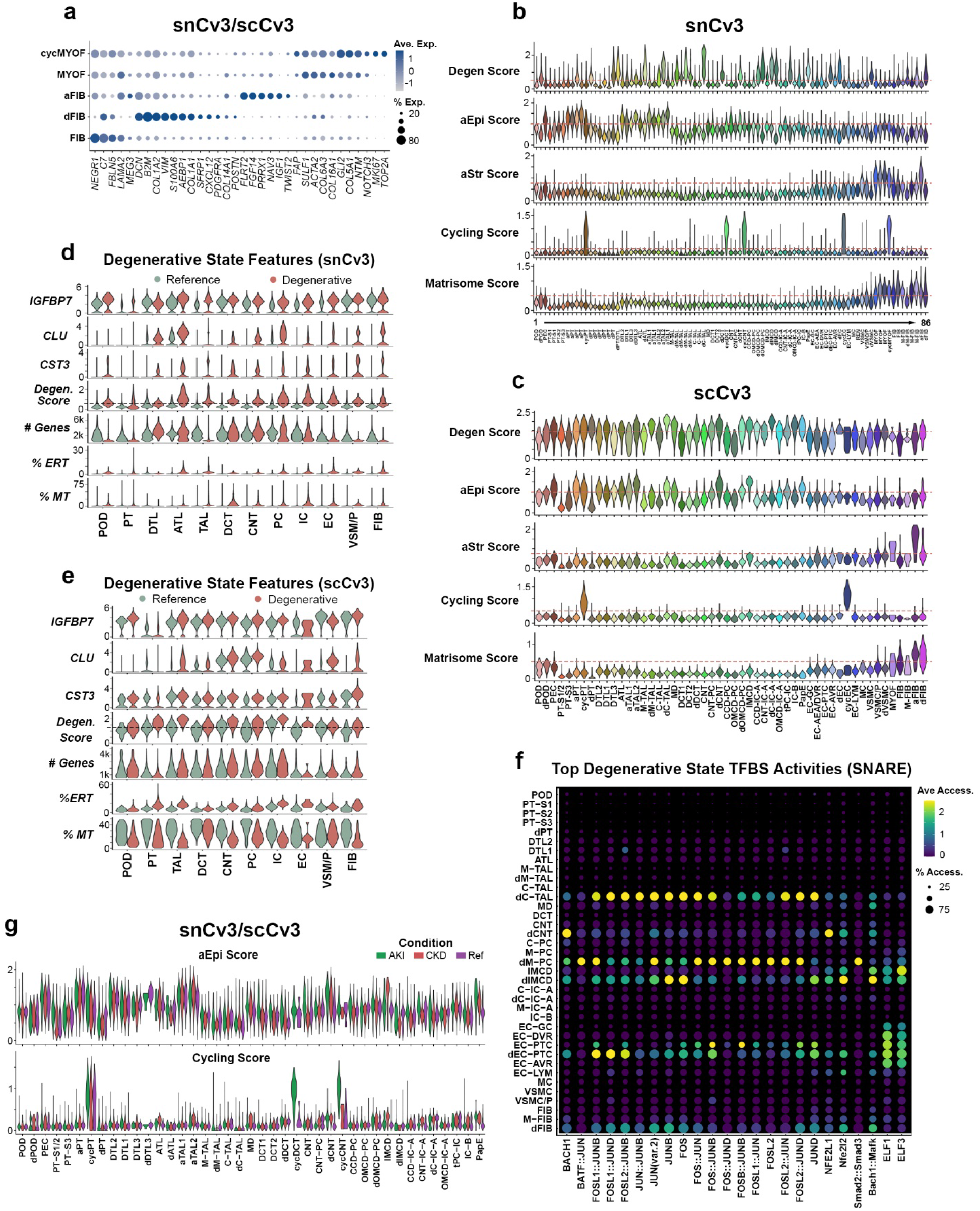
Single cell or nucleus altered state scoring. **a.** Dot plot showing averaged marker gene expression values (log scale) and proportion expressed for integrated snCv3/scCv3 reference, degenerative and adaptive stromal clusters. **b.** Violin plots showing adaptive state scores and ECM (matrisome) scores for snCv3 clusters. **c.** Violin plots as in (**b**) for scCv3 subclasses. **d.** Violin plots showing degenerative state scores and degenerative features (percent mitochondrial transcripts; percent ER or ribosomal transcripts; *CST3*, *CLU* and *IGFBP7* expression) for reference or degenerative states of snCv3 level 1 subclasses. **e.** Violin plots as in (**d**) for scCv3 level 1 subclasses. **f.** Dot plots showing SNARE2 average accessibilities (chromVAR) and proportion accessible for common degenerative TFBSs showing differential activity in 3 or more subclass level 1 cell types. **g.** Violin plots showing adaptive epithelial (aEpi) and cycling state scores for integrated snCv3/scCv3 level 3 subclasses split by condition (reference, AKI, CKD).

**Extended Data Figure 10.**
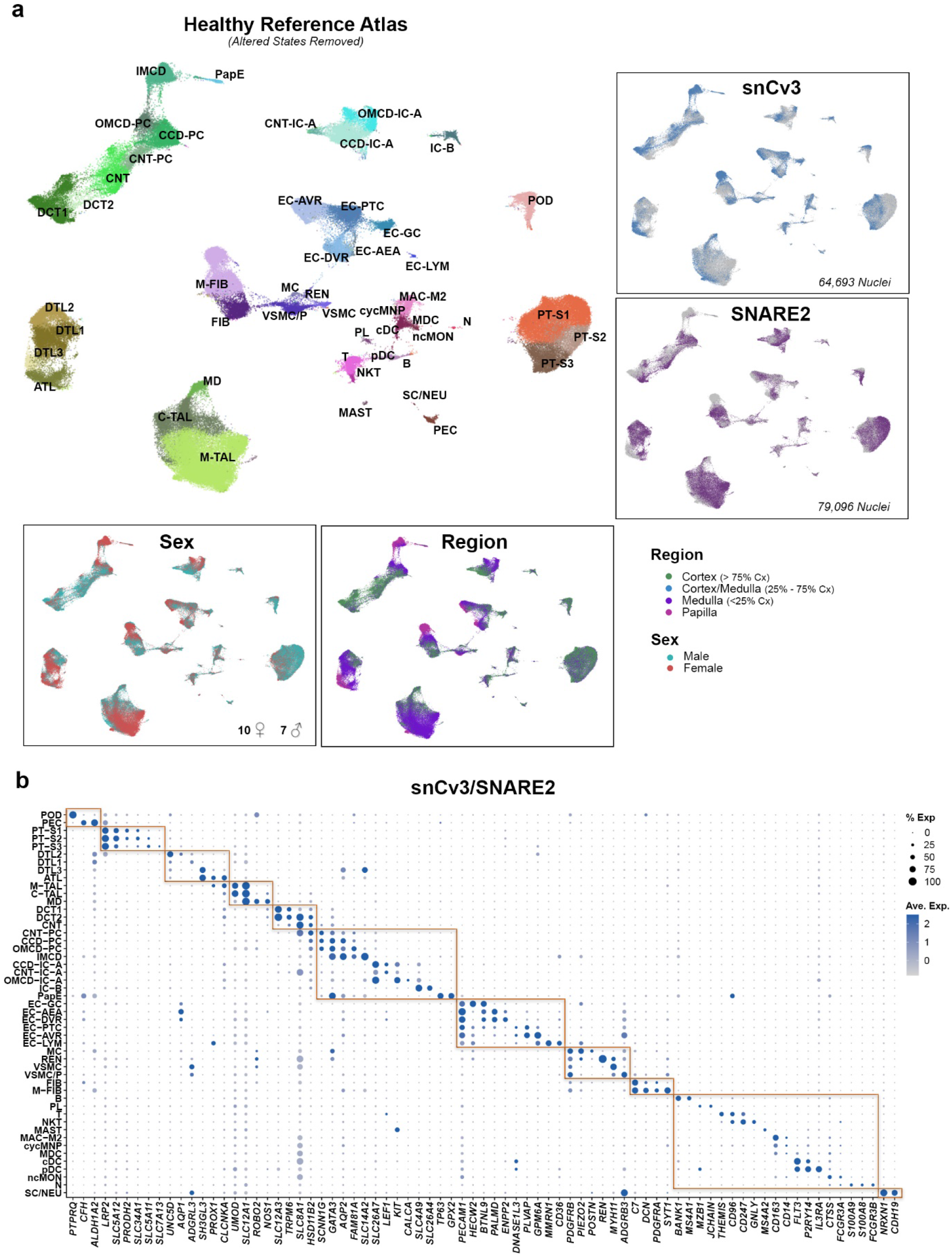
A healthy kidney reference atlas. **a.** UMAP plot of reference state level 3 subclasses for both snCv3 and SNARE2 (RNA) data. Insets show mapping of the tissue region, sex and assay identities. **b.** Dot plot showing averaged marker gene expression values (log scale) and proportion expressed for integrated snCv3/SNARE2 level 3 subclasses.

**Extended Data Figure 11.**
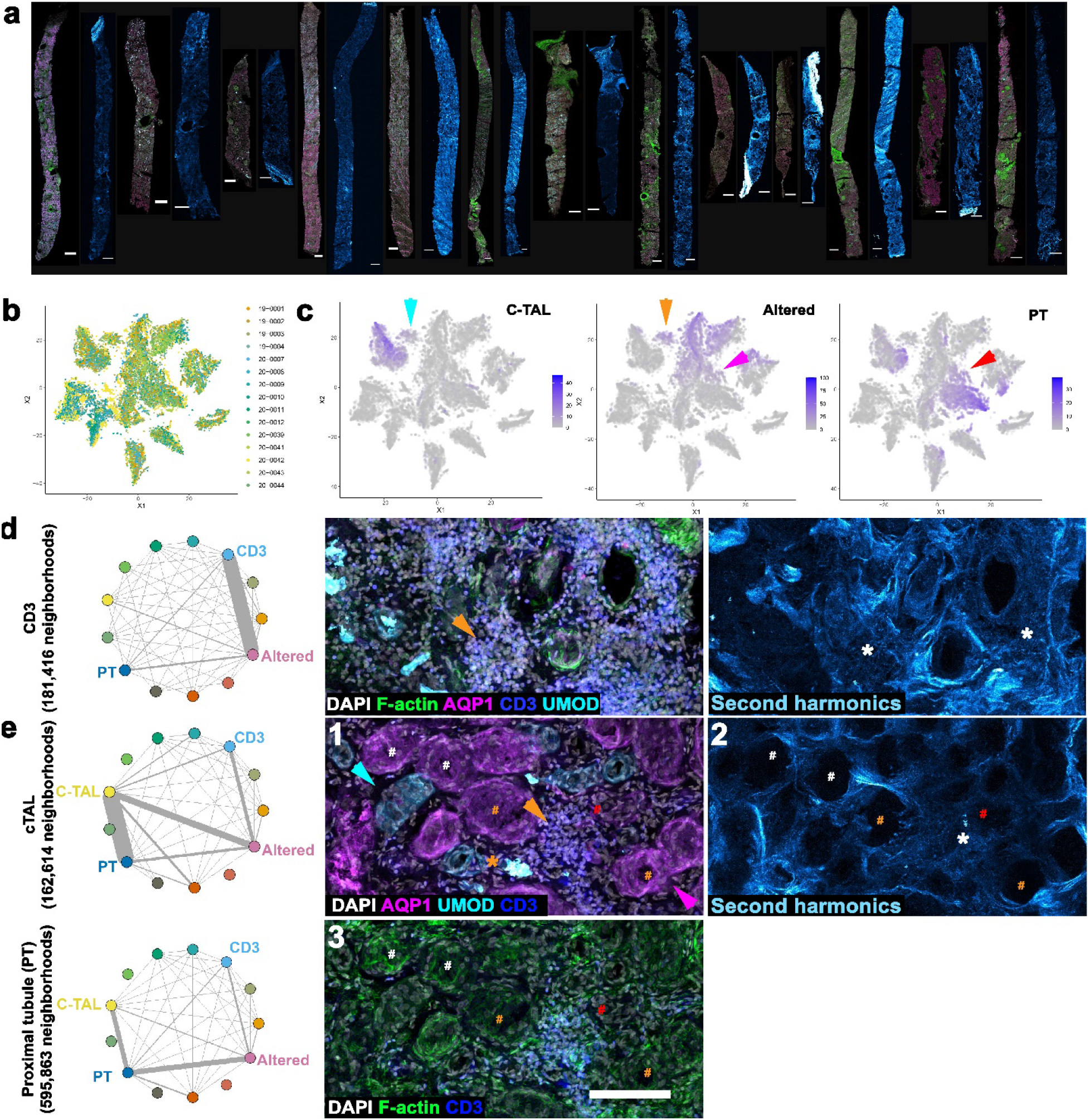
3D imaging identifies injury neighborhoods. **a**. Maximum intensity projections of immunofluorescence and second harmonic images for 13 example biopsies, scale bars 500 um. **b**. distribution of neighborhoods by specimen in neighborhood clusters plotted in tSNE space from **Fig. 4**. **c**. Feature plots of the number of cells per neighborhood for cortical TAL (C-TAL), altered morphology and proximal tubule (PT). C-TALs and PTs are found in neighborhoods with altered morphology, cyan and orange vs. red and magenta arrowheads. **d** and **e,** pairwise subset analysis of CD3+, PT and TAL (orange, magenta and cyan arrows respectively). CD3+ cells cluster in regions of fibrosis (orange arrowhead and white asterisks). UMOD positive casts associate with regions of injury and CD3+ cells (orange asterisk), the tubular epithelium is intact with brush borders (white #), has evidence of epithelial simplification (orange #) and shows a loss of marker and epithelial simplification (red #). Scale bar 100 um.

**Extended Data Figure 12.**
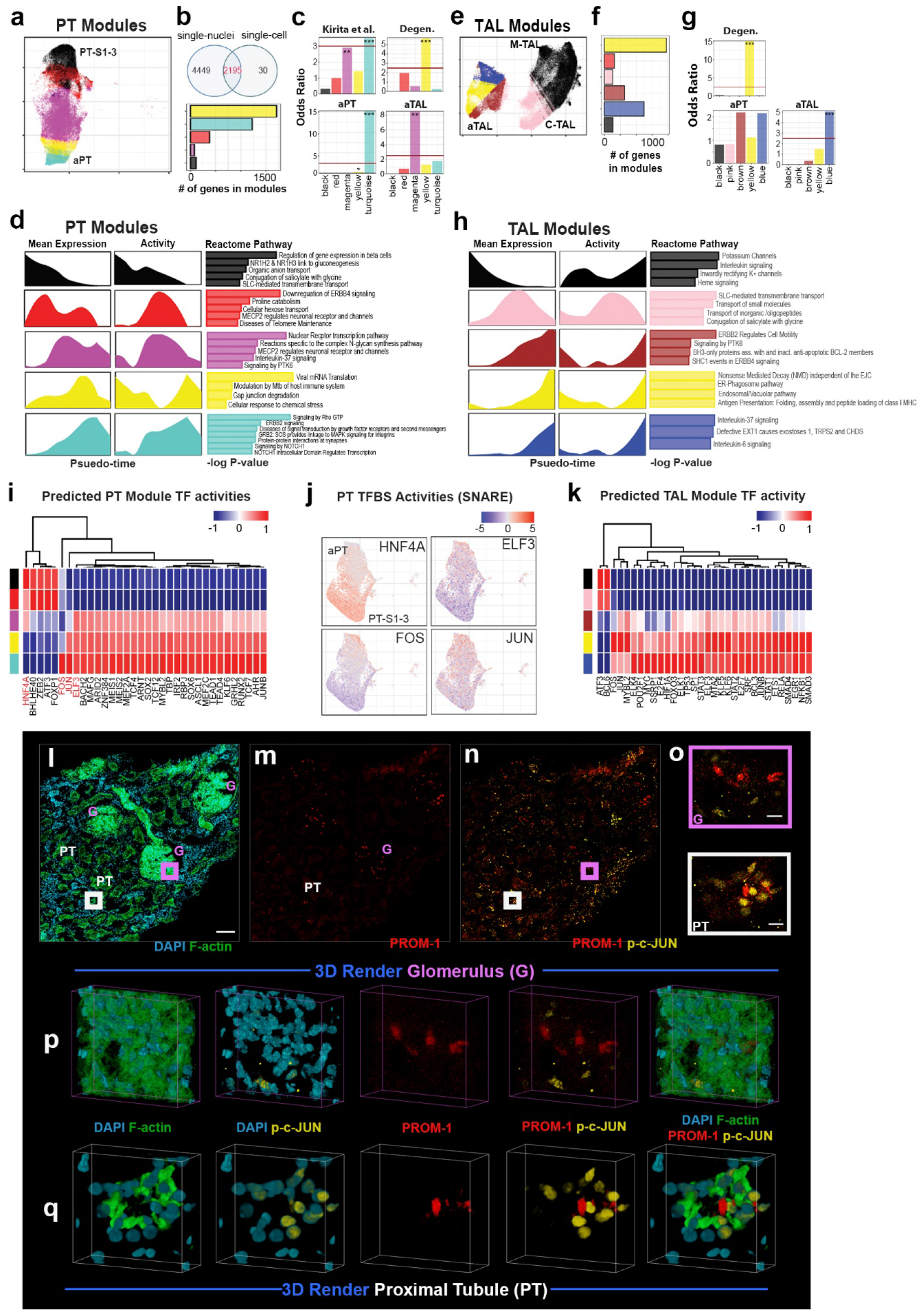
Expression signatures of adaptive epithelia. **a.** Umap embedding of PT cells colored by assigned modules (**Fig. 5**). **b.** Top: overlap of module associated genes in snCv3 and scCv3. Bottom: The number of genes in each PT module. **c.** Enrichment of failed to repair genes identified in Kirita et al.^3^ and genesets used for clinical outcome association (**Supplementary Table 27**) in each module (PT cells) identified by log-ratio test. **d.** The mean gene expression profile as a function of pseudo-time in PT modules and the top metabolic pathways in each identified module. **e.** Umap embedding of TAL cells colored by assigned modules (**Fig. 5**). **f.** The number of genes in TAL modules. **g.** Enrichment of genesets used for clinical outcome association (**Supplementary Table 27**) in each module (TAL cells) identified by log-ratio test. **h.** The mean gene expression profile as a function of pseudo-time in TAL modules and the corresponding top metabolic pathways in each identified module. **i, k.** Predicted TF transcription activities for cells in PT and TAL modules. **j.** Transcription binding site activities identified by SNARE2 for selected genes. **l-n.** 3D confocal imaging of a reference kidney tissue section stained for PROM-1 (red), Phopho-c-Jun (p-c-JUN, yellow), F-actin (with FITC phalloidin, green) and DNA with DAPI (cyan) (scale bar 100um). Regions of PROM-1 within a glomerulus (G) and a proximal tubule (PT) are marked with the magenta and white box, respectively and enlarged in (**o**) (scale bar 10um). **p.** and **q.** are snapshots of rendered 3D volumes V from the areas shown in (**o**). These areas show the association of PROM-1 expression with p-c-Jun+ cells in the tubules but not in glomerular cells. 3D rendering was performed using the Voxx software from the Indiana Center for Biological Microscopy (voxx.sitehost.iu.edu/).

**Extended Data Figure 13.**
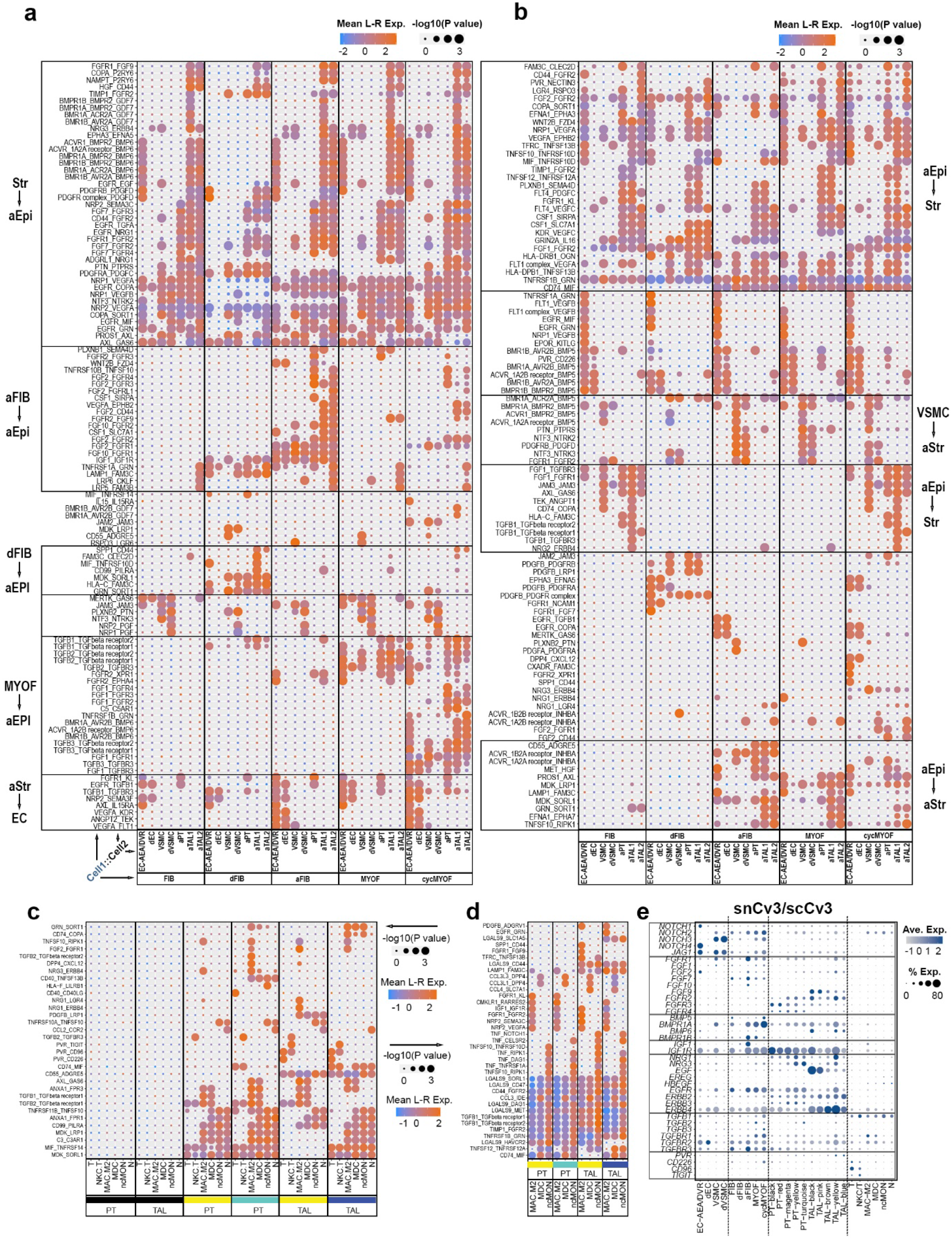
Ligand receptor signaling in the fibrotic niche. **a.** Significant (p value < 0.05) secreted ligand and receptor interactions (excluding integrins) identified for signaling from the stroma to vascular and adaptive epithelial cells. **b.** Significant (p value < 0.05) secreted ligand and receptor interactions (excluding integrins) identified for signaling from vascular and adaptive epithelial cells to the stroma. **c.** Significant (p value < 0.05) secreted ligand and receptor interactions (excluding integrins) identified for signaling from adaptive epithelial trajectory modules to immune cells. Only interactions that were also not significant (p value > 0.05) in reference modules were plotted. **d.** Significant (p value < 0.05) secreted ligand and receptor interactions (excluding integrins) identified for signaling from macrophage-type immune cells to the adaptive epithelial modules. **e.** Dot plot showing averaged gene expression values (log scale) and proportion expressed for select ligands and receptors. All ligand-receptor analyses and expression plots were for integrated snCv3/scCv3 level 3 subclasses or modules.

**Extended Data Figure 14.**
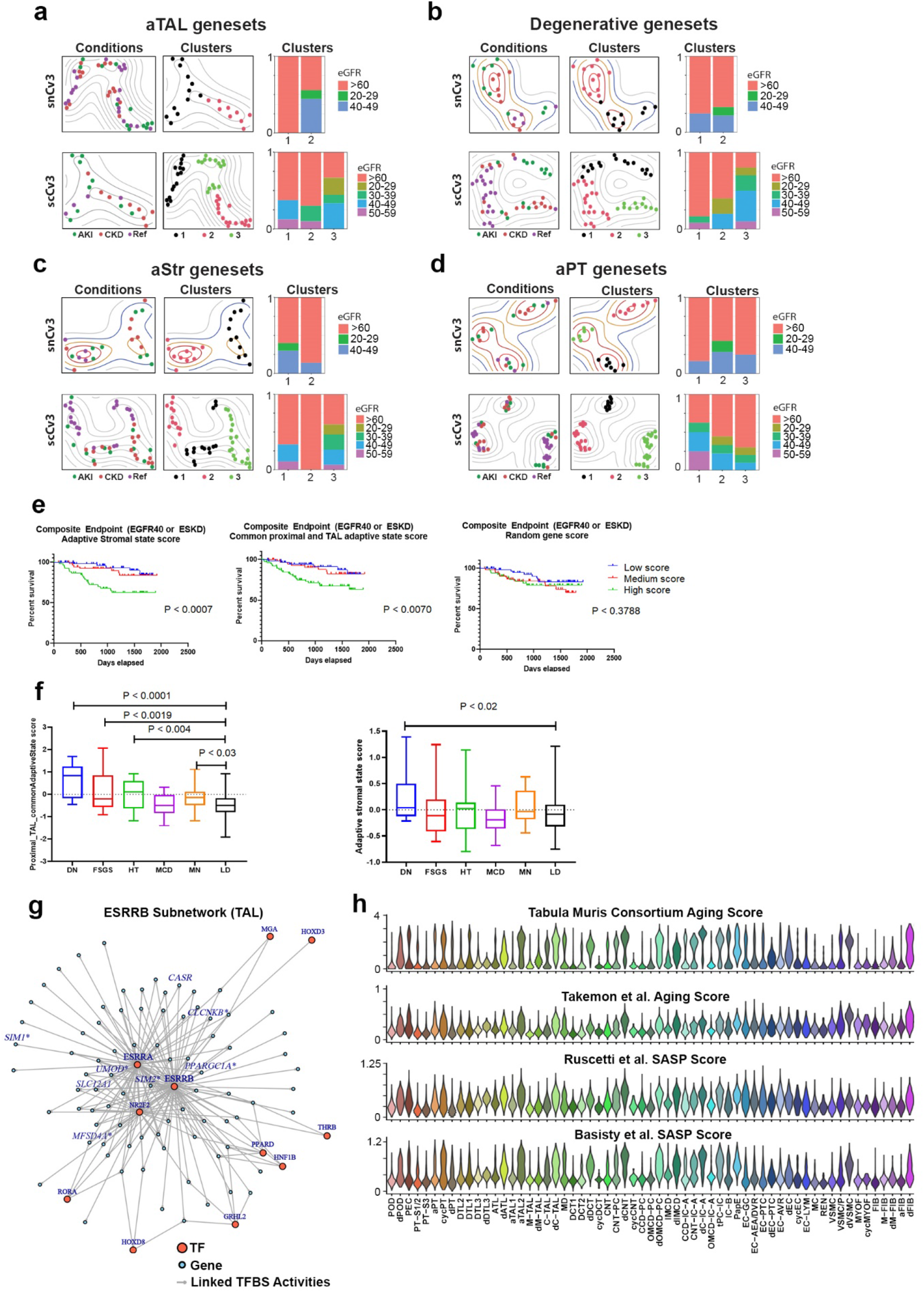
Association of cell state scores with clinical phenotypes. **a.** Left panels: grouping of patient-level expression profiles for the aTAL geneset used for clinical outcome association (**Supplementary Table 27**) for snCv3 (Top) and scCv3 (Bottom). Right panels: the distribution of eGFR among the identified groups. **b.** Plots as in (**a**) for the degenerative geneset used for clinical outcome association. **c.** Plots as in (a) for the aStr geneset used for clinical outcome association. **d.** Plots as in (**a**) for the aPT geneset used for clinical outcome association. **e.** Unadjusted Kaplan Meier curves by aStr and common aPT and aTAL state scores for composite of ESRD or 40% drop in eGFR from time of biopsy in Neptune adult patient cohort. P values from log-rank tests for trend are shown. A score generated using 100 randomly selected genes failed to show any correlation with disease survival. **f.** Boxplot of aStr and common aPT and aTAL cell state scores by kidney disease groups in the ERCB cohort. Significant P values from unpaired t-tests between disease groups and LD are shown. The DN patient group had significantly higher aStr and common aPT and aTAL cell state scores compared to LD. **g.** ESRRB subnetwork of TF connections to target genes generated using SNARE2 RNA and AC data, demonstrating a central role for ESRRB in regulating TAL marker genes. Inset shows the ESRRB motif. Boxes represent ESRRB target genes showing causal variant enrichment within linked regulatory regions (AC peaks). **h.** Violin plots show gene expression scores for gene sets associated with aging (Tabula Muris Consortium^46^ and Takemon et al.^63^) or SASP (Ruscetti et al.^64^ or Basisty et al.^65^).

## Supplementary Tables

**Supplementary Table 1.** Summary of omic experiments

**Supplementary Table 2.** Summary of 3D imaging and spatial transcriptomic experiments

**Supplementary Table 3.** Sample Clinical Data

**Supplementary Table 4.** snCv3 cluster annotations

**Supplementary Table 5.** snCv3 Cell Type Marker Genes

**Supplementary Table 6.** scCv3 cluster annotations

**Supplementary Table 7.** SNARE2 (RNA/AC) cluster annotations (Post-AC processing and clusters > 50 nuclei, Methods)

**Supplementary Table 8.** snCv3 NSForestv2 Computational Marker Genes

**Supplementary Table 9.** scCv3 NSForestv2 Computational Marker Genes

**Supplementary Table 10.** Differentially Accessible Regions (DARs) associated with subclass level marker genes

**Supplementary Table 11.** Motif enrichments for marker gene associated Differentially Accessible Regions (DARs)

**Supplementary Table 12.** Cell type specific TFBS activities

**Supplementary Table 13.** Cell state definitions

**Supplementary Table 14.** aEpi Differentially expressed genes (snCv3)

**Supplementary Table 15.** Conserved degenerative state marker genes

**Supplementary Table 16.** Conserved adaptive epithelial (aEpi) state marker genes

**Supplementary Table 17.** Conserved adaptive stromal (aStr) state

**Supplementary Table 18.** Conserved cycling state marker genes

**Supplementary Table 19.** Adaptive PT trajectory gene expression modules (snCv3)

**Supplementary Table 20.** Adaptive TAL trajectory gene expression modules (snCv3)

**Supplementary Table 21.** Adaptive epithelial module reactome pathway analyses

**Supplementary Table 22.** TF activities predicted from aEpi gene modules

**Supplementary Table 23.** TF activities identified for aEpi trajectory modules using SNARE2

**Supplementary Table 24.** CellPhoneDB significant (p value < 0.05) ligand-receptor pairs (mean expression values) between adaptive state subclasses and interstitial/vascular cell types

**Supplementary Table 25.** CellPhoneDB significant (p value < 0.05) ligand-receptor pairs (mean expression values) between adaptive modules and interstitial/vascular cell types

**Supplementary Table 26.** CellPhoneDB significant (p value < 0.05) ligand-receptor pairs (mean expression values) used in circos plots

**Supplementary Table 27.** Altered state gene sets used for clinical outcomes assessment

**Supplementary Table 28.** Motif enrichments within causal variant SNP peaks associated with the TAL

**Supplementary Table 29.** Genesets that distinguish AKI and CKD patients in aPT and aTAL trajectories

**Supplementary Table 30.** Time of biopsy characteristics of participants in Neptune cohort used in this study.

**Supplementary Table 31.** Antibodies and dilutions used in mesoscale 3D imaging for 3D cytometry

**Supplementary Table 32.** Antibodies and dilutions used in confocal imaging

**Supplementary Table 33.** Tabulation of select marker based 3D cytometry results

## Notes

https://portal.hubmapconsortium.org/

https://atlas.kpmp.org

https://azimuth.hubmapconsortium.org/

### Competing Interest Statement

The authors have declared no competing interest.

